# Activity-based urinary biomarkers of response and resistance to checkpoint blockade immunotherapy

**DOI:** 10.1101/2020.12.10.420265

**Authors:** Quoc D. Mac, Congmin Xu, James R. Bowen, Anirudh Sivakumar, Hathaichanok Phuengkham, Fang-Yi Su, Samuel Z. Stentz, Hyoungjun Sim, Adrian M. Harris, Tonia T. Li, Peng Qiu, Gabriel A. Kwong

## Abstract

Immune checkpoint blockade (ICB) therapy has transformed cancer treatment, yet most patients do not derive clinical benefit and responders can acquire resistance to therapy. Noninvasive biomarkers are needed to indicate early on-treatment response and resistance mechanisms. Here we developed <u>I</u>mmu<u>N</u>e <u>S</u>ensors for mon<u>I</u>torinG c<u>H</u>eckpoint blockade <u>T</u>herapy (INSIGHT), which comprises a library of mass-barcoded peptide substrates conjugated to αPD1 antibodies, as therapeutic sensors. Following systemic administration, INSIGHT carries out the dual role of reinvigorating T cell function and profiling T cell and tumor proteases by the release of cleaved peptides into urine for noninvasive detection by mass spectrometry. We show that an αPD1 therapeutic sensor for Granzyme B discriminates early treatment responses before tumor volumes significantly diverge from isotype controls in murine models of colorectal cancer. To differentiate mechanisms of resistance by multivariate analysis, we design a mass-barcoded, 14-plex INSIGHT library to profile proteases differentially expressed by tumors harboring B2m or Jak1 loss-of-function mutations. We find that binary classifiers trained on urine samples indicate response to αPD-1 therapy as early as the start of the second dose, and discriminate B2m from Jak1 resistance with high sensitivity and specificity (AUROCs > 0.9). Our data supports the use of activity-based biomarkers for early on-treatment response assessment and classification of refractory tumors based on resistance mechanisms.

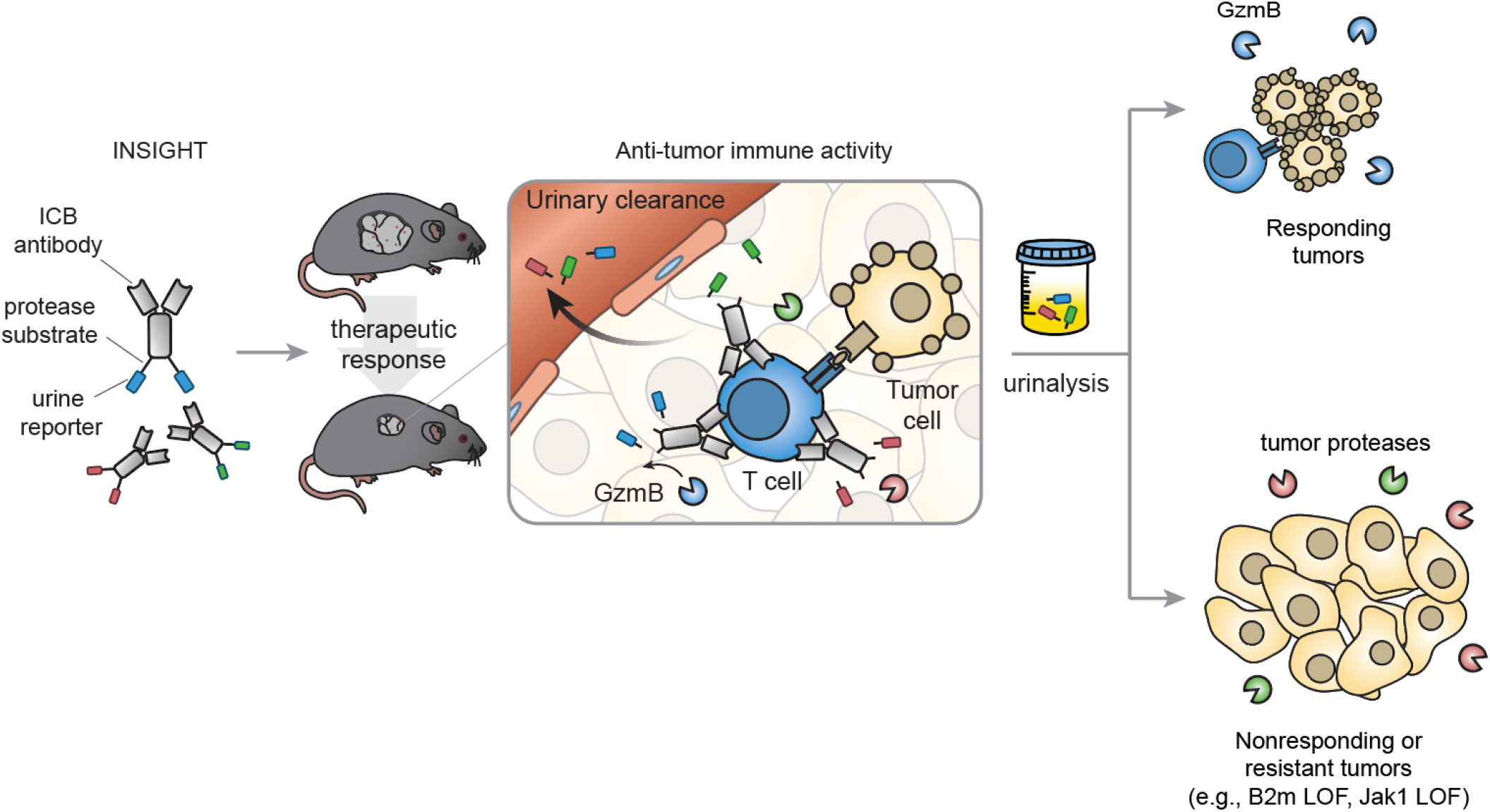

## Introduction

Immune checkpoint blockade (ICB) therapy has transformed the treatment of cancer for patients across a broad range of malignancies^1,2^. ICB involves the administration of antibodies that block inhibitory checkpoint molecules, such as the cytotoxic T lymphocyte-associated protein 4 (CTLA-4) or the programmed cell death protein 1 (PD-1), to reinvigorate an anti-tumor T cell response. Despite the potential for ICB to produce durable clinical outcomes, a large fraction of patients do not derive clinical benefit^1,3^. Objective response rates remain below ~25% in many cancer types, largely due to immunosuppressive factors in the tumor microenvironment (TME) (e.g., Tregs or MDSCs) and primary tumor-intrinsic mutations^1^. In addition, responsive tumors can acquire resistance during therapy such as in metastatic melanoma where up to one-third of patients with initial responses to ICB therapy eventually relapse^3^. Both primary and acquired resistance are driven by mechanisms that enable tumor cells to evade anti-tumor immune responses, including defects in antigen presentation or IFNγ response pathway^3,4^. Therefore, developing noninvasive biomarkers of immune response and resistance to ICB has emerged as a clinical priority^5^.

Patient responses to ICB therapy are currently assessed using a combination of radiographic, tumor, and serum biomarkers^5^. Radiographic evaluation by RECIST criteria is the standard assessment method and occurs after the first cycle of ICB therapy, which consists of 3-4 doses administered within an 8-12-week window^6–8^. The observation of atypical patterns of response to ICB has motivated continual refinement to the timing and frequency of radiographic assessment such as the development of immune-related response criteria (e.g., irRC, irRECIST) to account for phenomenon like pseudoprogression^5,9^. Tumor biomarkers such as PD-L1 expression have been shown to enrich for populations with clinical benefit, but have limitations as predictive biomarkers as at least ~40-50% patient tumors with PD-L1 positivity do not experience objective responses^5,10^. Other tumor biomarker strategies, such as assessing on-treatment changes in tumor mutational burden (TMB) by whole exome sequencing^11^, are promising and have been found to correlate with αPD1 response. However, these approaches require serial biopsies, which in practice are not typically collected over the course of therapy with attendant patient risks. Therefore, considerable interest is focused on identifying noninvasive biomarkers to allow longitudinal and quantitative assessment. These include quantifying changes in T cell clonality or circulating tumor DNA (ctDNA) levels, which have been shown to be detectable within 3-4 weeks of treatment and correlate with objective response and overall survival^12–14^. These studies highlight the considerable interest and need for noninvasive and longitudinal assessment strategies to track response and resistance to ICB therapy early on-treatment.

Proteases play fundamental roles in cancer biology, immunity, and anti-tumor responses and therefore may provide a new mechanism to evaluate ICB therapy. Tumor-dysregulated proteases (e.g., matrix metalloproteases, cathepsins) are involved in proteolytic cascades that modify the tumor microenvironment (TME) during angiogenesis, growth, and metastasis^15,16^. In addition, T cell-mediated tumor control is primarily carried out by granzymes, which are serine proteases, released by cytotoxic T cells^17^. The ubiquity of protease dysregulation has motivated the development of molecular imaging probes for visualizing tumor or T cell proteases^18–21^, as well as synthetic biomarkers for multiplexed quantification of protease activity from urine^22–27^. Building on these studies, we developed Immune Sensors for monItorinG cHeckpoint blockade Therapy (INSIGHT) to detect tumor and immune proteases during treatment as activity-based biomarkers of response and resistance. INSIGHT immune sensors consist of mass-barcoded protease substrates conjugated to ICB antibodies that during the course of treatment are cleaved by proteases, triggering the release of reporters that filter into urine. After urine collection, cleaved reporters are quantified by mass spectrometry according to their mass barcode. In preclinical animal models, we show that binary classifiers trained on protease signatures by machine learning indicate on-treatment responses as early as the start of the second dose and differentiate B2m and Jak1 resistance with high sensitivity and specificity.

## Results

### Antibody-peptide therapeutic sensors retain target binding and in vivo therapeutic efficacy

We first characterized target binding and therapeutic efficacy of ICB antibody-peptide conjugates. As a representative formulation, we coupled a fluorescently labeled peptide substrate selective for murine GzmB (IEFDSG^26^) to αPD1 (clone 8H3) to form an αPD1-GzmB sensor conjugate (αPD1-GS) (**Fig. 1a**). To determine whether peptide conjugation would interfere with PD1 binding, we tested different peptide:antibody stoichiometric ratios (0, 1, 3, 5, 7) and quantified binding to recombinant PD1 by ELISA. We observed negligible differences in EC50 at a 1:1 ratio compared to unmodified αPD1 (3.6 vs. 2.1 nM respectively) (**Fig. 1b**) but at higher ratios, a gradual reduction in binding (up to 24 nM at a 7:1 ratio) (**SFig. 1a**). To confirm that these results were not clone dependent, we coupled GzmB peptides to another αPD1 clone (29F.1A12) at a 1:1 ratio and found that target binding was likewise preserved between αPD1-GS and unconjugated antibody (EC50 = 0.15 nM vs. 0.18 nM) (**Fig. 1c**). Based on these results, we used a 1:1 conjugation ratio for all subsequent studies.

**Figure 1.**
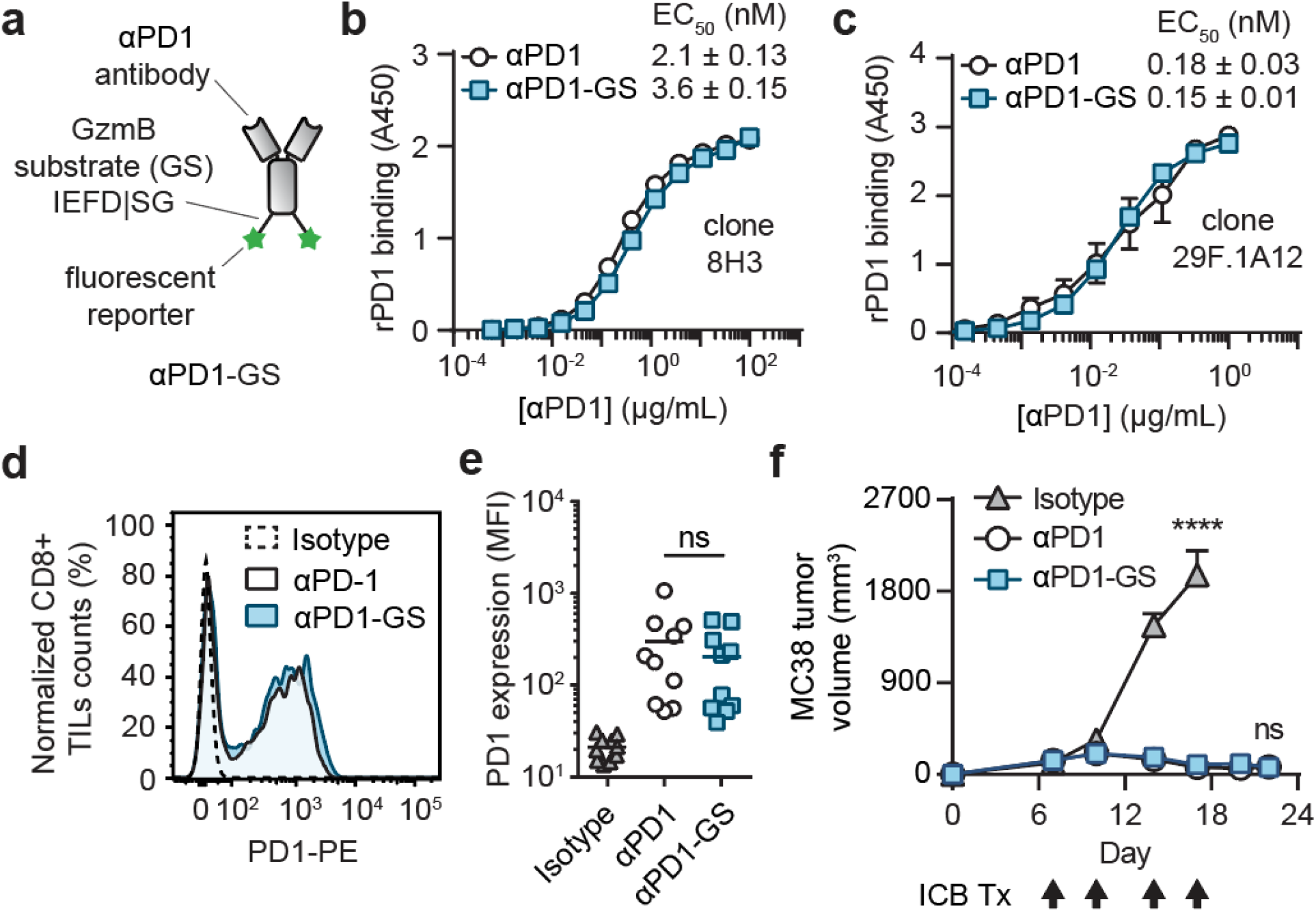
Antibody binding and therapeutic efficacy are unaffected by peptide conjugation. **a**, αPD1-GzmB sensor conjugates (αPD1-GS) consist of αPD1 therapeutic antibody decorated with reporter-labeled GzmB peptide substrates (GS; AA sequence: IEFDSG). **b**, ELISA assays comparing binding affinity of αPD1-GS with unconjugated αPD1 using the mouse αPD1 clone 8H3 (log(agonist) vs. normalized response fitting function, n = 3). **c**, ELISA assays comparing binding affinity of αPD1-GS with unconjugated αPD1 using the rat αPD1 clone 29F.1A12 (log(agonist) vs. normalized response fitting function, n = 3). **d**, Representative flow cytometry histogram showing PD-1 expression of CD8+ TILs isolated from MC38 tumors. The same sample was divided and stained with either αPD1-GS, αPD1, or IgG1 isotype control. **e**, Quantified plot of PD-1 expression showing the median fluorescence intensity (MFI) of samples stained with either αPD1-GS, αPD1, or IgG1 isotype control (one-way ANOVA with Tukey’s post-test and correction for multiple comparisons, ns = not significant, n = 10). **f**, Tumor growth curves of MC38 tumors treated with αPD1-GS, αPD1, or IgG1 isotype control (two-way ANOVA with Tukey’s post-test and correction for multiple comparisons, ****P < 0.0001, n = 6).

We next evaluated target binding of αPD1-GS to tumor infiltrating lymphocytes (TILs) isolated from MC38 tumors since ligand presentation of plate-bound recombinant PD1 may differ from endogenous PD1 expressed by T cells. We used the MC38 colon adenocarcinoma syngeneic tumor model because these cancer cells have a high mutation burden, which has been shown to lead to an endogenous T cell infiltrate following αPD1 monotherapy^28^. Flow cytometry analysis of CD8+ TILs stained with either αPD1-GS or unmodified αPD1 showed statistically equivalent PD1 expression by median fluorescence intensity (MFI), indicating that peptide conjugation did not significantly affect target binding to endogenous PD1 expressed on cell surfaces (n = 10, **Fig. 1d, e**). We further confirmed that peptide conjugation did not affect therapeutic efficacy by comparing anti-tumor responses. Following a treatment schedule that involved four doses of antibody to C57BL/6 mice bearing MC38 tumors, we observed no statistical difference in tumor burden in mice given αPD1-GS or unmodified αPD1. Both formulations resulted in smaller tumors that were statistically significant compared to animals given IgG1 isotype control (P ≤ 0.0001, n = 6, **Fig. 1f**). Taken together, these data demonstrate that coupling peptides at a low molar ratio to αPD1 does not affect target binding or *in vivo* therapeutic efficacy.

### αPD1-GS detects GzmB activity during T cell killing of tumor cells

We next tested the ability of αPD1-GS to monitor GzmB activity in a T cell killing assay. To quantify cleavage activity by fluorimetry, we coupled GzmB peptides containing a fluorophore-quencher pair (5FAM-AIEFDSG-CPQ2) to αPD1. (**Fig. 2a**). We assessed substrate specificity by incubating αPD1-GS with fresh mouse serum, tumor-associated proteases (e.g., cathepsin B, MMP9), or coagulation and complement proteases (e.g., C1s, thrombin). While incubation with recombinant GzmB led to a rapid increase in sample fluorescence, incubation with mouse serum or recombinant proteases did not result in detectable increases in fluorescence that would indicate cross-cutting of our sensors (**Fig. 2b**). To evaluate αPD1-GS activation in the context of a T cell killing assay, we cocultured Pmel T cells with gp100-expressing B16 melanoma cells at increasing effector to target cell ratios (0, 1, 5, 10) and verified statistically significant increases in both supernatant GzmB by ELISA and target cell death by lactose dehydrogenase (LDH) release (n = 3, **Fig. 2c, d**). Under these co-culture conditions, we observed significant increases in fluorescence only in cocultures incubated with αPD1-GS, but not in control wells containing unmodified αPD1 antibody or αPD1 conjugated with a control peptide substrate (5FAM-ALQRIYK-CPQ2) (n = 3, **Fig. 2e**). We also did not observe αPD1-GS activation in cocultures of OT1 T cells and B16 cancer cells, which do not express the OVA antigen. (P ≤ 0.0001, n = 4, **Fig. 2f**). Collectively, these data demonstrate that αPD1-GS is selectively cleaved by GzmB and can be used to detect T cell killing of tumor cells.

**Figure 2.**
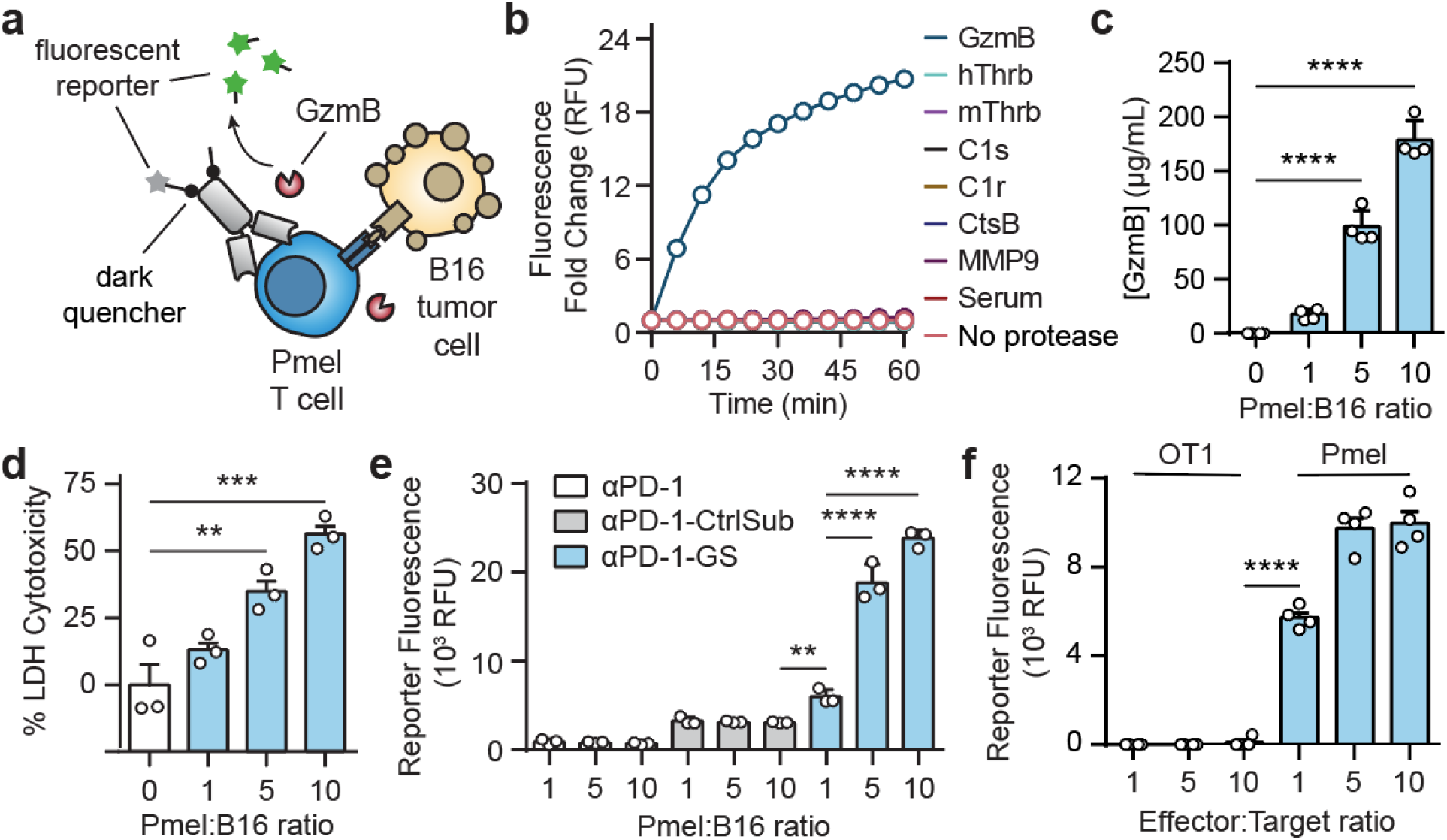
Sensing T cell killing of tumor cells by antibody-GzmB sensor conjugates. **a**, αPD1 antibody was conjugated with fluorescently-quenched peptide substrates for GzmB. Upon incubating these conjugates with transgenic Pmel T cells and B16 tumor cells, secreted GzmB cleaved peptide substrates, separating the fluorescent reporter from the internal quencher and resulting in an increase in sample fluorescence. **b**, *In vitro* protease cleavage assays showing normalized fluorescence of αPD1-GS after incubation with recombinant GzmB (blue), mouse serum (red), and other bystander proteases (n = 3). **c**, ELISA quantification of GzmB from T cell killing assays in which Pmel T cells were incubated with B16 target cells at different T cell to target cell ratios (one-way ANOVA with Dunnett’s post-test and correction for multiple comparisons, ****P < 0.0001, n = 4). **d**, Bar plot quantifying percent of cell cytotoxicity as measured by LDH assay from cocultures of Pmel T cells with B16 target cells (one-way ANOVA with Dunnett’s post-test and correction for multiple comparisons, ***P < 0.001, n = 3). **e**, Activity assays showing sample fluorescence after incubating αPD1-GS, αPD1, and an αPD1 conjugate with control substrates (αPD1-CtrlSub) with cocultures of Pmel T cells with B16 target cells (two-way ANOVA with Tukey’s post test and correction for multiple comparisons, ****P < 0.0001, n = 3). **f**, Activity assays showing sample fluorescence after incubating αPD1-GS with cocultures of Pmel or OT1 transgenic T cells with B16 target cells (two-way ANOVA with Tukey’s post test and correction for multiple comparisons, ****P < 0.0001, n = 3).

### Noninvasive detection of early on-treatment response to ICB therapy

We next evaluated the potential of αPD1-GS to noninvasively detect response to treatment in mouse models based on GzmB activity alone. Because free peptides can be rapidly degraded in blood but have improved pharmacokinetic profiles when conjugated to an antibody or protein scaffold^29,30^, we first quantified the plasma concentration of uncleaved αPD1-GS following intravenous administration to determine peptide stability. We developed an indirect ELISA that uses plate-bound PD1 to capture αPD1-GS and a detection antibody specific for the FAM reporter at the termini of the peptide substrate to differentiate between cleaved and uncleaved conjugates (**SFig. 2a**). In validation assays, we compared ELISA signals from samples that contained αPD1-GS with or without preincubation with recombinant GzmB. Whereas αPD1-GS was readily detected compared to unmodified αPD1, we observed dose dependent reduction in signals for αPD1-GS samples treated with GzmB (n = 3, **SFig. 2b, 2c**), validating the ability to discriminate between cleaved and uncleaved conjugates. Using this assay, we determined that the circulation half-life of uncleaved αPD1-GS was several hours and statistically equivalent to unmodified αPD1 antibody (3.9 ± 1.3 h vs 6.5 ± 4.2 h, n = 3, two-way ANOVA) (**Fig. 3a**), indicating peptide stability in circulation.

**Fig 3.**
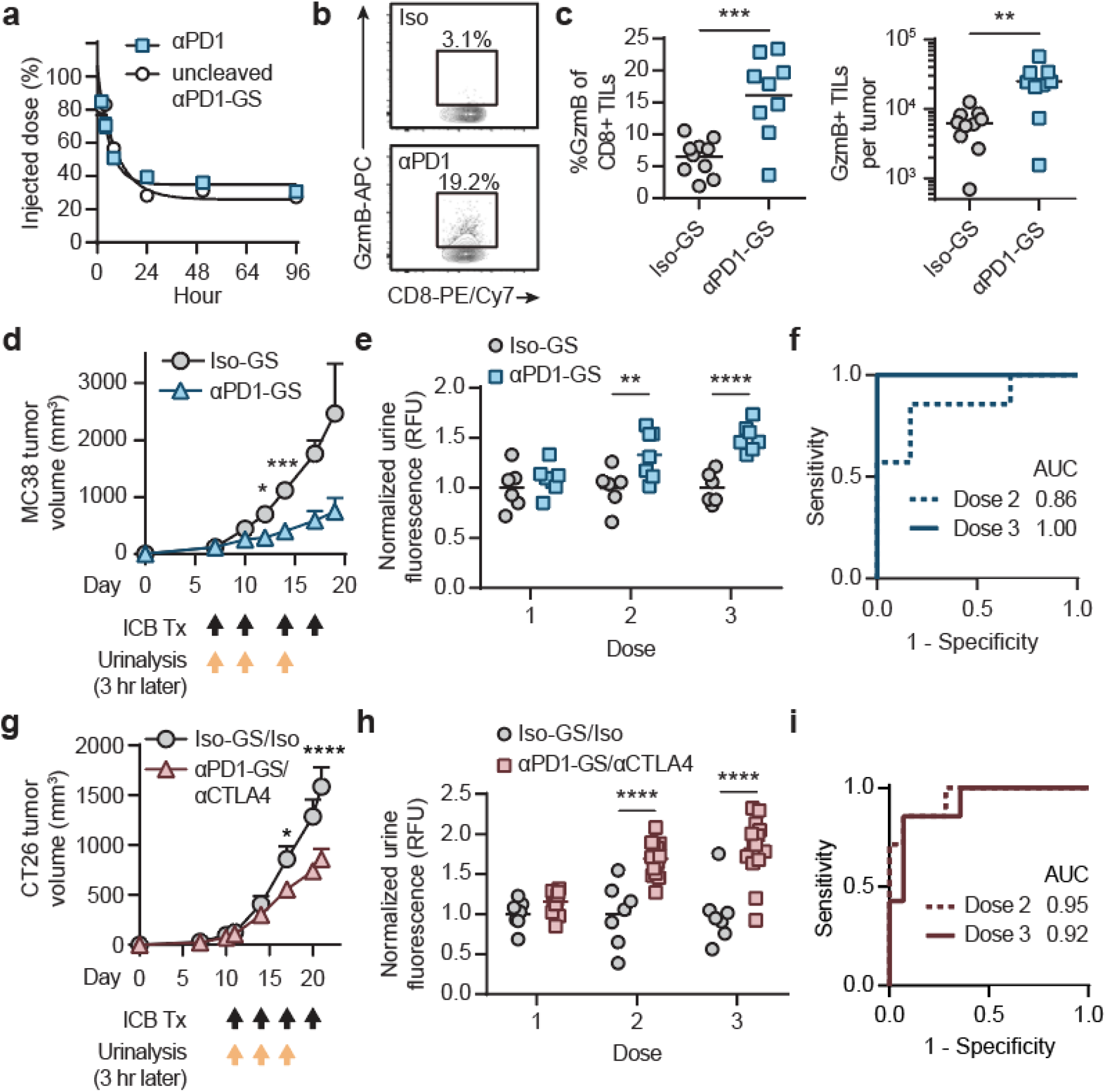
Urinary detection of ICB therapeutic response by administration of antibody-GzmB sensor conjugates. **a**, Half-life measurements of intact αPD1-GS and unconjugated αPD1 antibody (one phase decay fitting function, n = 3). **b**, Representative flow cytometry plots showing intracellular GzmB expression of CD8+ TILs from MC38 tumors treated with either αPD1-GS or IgG1 isotype antibody conjugated with the GzmB peptide substrates (Iso-GS). **c**, Quantified plots showing percentages of GzmB+ cells within the CD8+ TILs or the numbers of GzmB+CD8+ TILs that were isolated from MC38 tumors treated with either αPD1-GS or Iso-GS (two-sided Student’s t-test, n = 9-10). **d**, Tumor growth curves of MC38 tumor bearing mice treated with either αPD1-GS or Iso-GS (two-way ANOVA with Sidak’s post test and correction for multiple comparisons, ***P < 0.001, n = 6-7). Black arrows denote the treatment time points. **e**, Left: normalized urine fluorescence of mice with MC38 tumors after each administration of αPD1-GS or Iso-GS (two-way ANOVA with Sidak’s post test and correction for multiple comparisons, ****P < 0.0001, n = 6-7). **f**, Receiver-operating-characteristic (ROC) analysis showing the diagnostic specificity and sensitivity in differentiating between mice treated with aPD1-GS vs. Iso-GS using urine signals on the second (AUC = 0.857, 95% CI = 0.643-1.00) or the third dose (AUC = 1.00, 95% CI = 1.00-1.00). **g**, Tumor growth curves of CT26 tumor bearing mice treated with combination therapy of αPD1-GS and αCTLA4 or combination of matched isotype controls (two-way ANOVA with Sidak’s post test and correction for multiple comparisons, ****P < 0.0001, n = 7-14). Black arrows denote the treatment time points. **h**, Normalized urine fluorescence of mice with CT26 tumors after each administration of αPD1-GS and αCTLA4 or matched isotype controls (two-way ANOVA with Sidak’s post test and correction for multiple comparisons, ****P < 0.0001, n = 7-14). **i**, ROC analysis showing the diagnostic specificity and sensitivity of αPD1-GS in differentiating between responders to ICB combination therapy from off-treatment controls using urine signals on the second (AUC = 0.949, 95% CI = 0.856-1.00) or the third dose (AUC = 0.92, 95% CI = 0.795-1.00)

We evaluated αPD1-GS to detect response in C57BL/6 mice bearing MC38 tumors. We confirmed significantly elevated expression of GzmB in CD8+ TILs following two doses of αPD1-GS compared to control mice that received an isotype antibody conjugated with the same peptide (Iso-GS) (P ≤ 0.001, n = 9, **Fig. 3b, c**). To evaluate the potential for serial on-treatment response assessment, we quantified the concentration of cleaved fluorescent reporters in urine samples that were collected within 3 hours after each dose was administered (day 7, 10, 14) (**Fig. 3d**). At the start of the first dose on day 7, urine signals from both cohorts of mice were statistically identical as expected. By contrast, urine signals were significantly elevated in mice treated with αPD1-GS at the start of the second dose on day 10 (P ≤ 0.01, n = 6-7) when tumors were statistically equivalent in volume compared to control mice that received Iso-GS (255 mm^3^ vs. 441 mm^3^, P = 0.68, n = 6-7). This difference in urine signals was further accentuated by the start of the third dose on day 14 (P ≤ 0.0001, n = 6-7) (**Fig. 3e**). Receiver operator characteristic (ROC) analysis of reporter levels in urine samples revealed an area under curve (AUC) of 0.86 and 1.00 for dose 2 and 3 respectively (**Fig. 3f**), indicating the ability to differentiate ICB response with high sensitivity and specificity.

We further sought to confirm urinary detection in a different preclinical model using BALB/c mice bearing syngeneic CT26 tumors that respond to combination therapy (αPD1 and αCTLA4) but minimally to monotherapy (αPD1 or αCTLA4)^31,32^. Compared to matched isotype control conjugates, monotherapy with either αPD1-GS or αCTLA4-GS did not result in statistical differences in tumor burden and urine signals across all doses (**SFig. 3a, b, c, d**). By contrast, combination treatment with αPD1-GS and αCTLA4 resulted in significantly lower tumor burden (P ≤ 0.0001, n = 7-14, **Fig. 3g**), higher levels of GzmB+ CD8+ TILs (P ≤ 0.05, n = 7, **SFig. 4a, b**), and significant increases in urine signals at the start of the second or third dose (AUROC = 0.95 and 0.92 respectively, **Fig. 3h**). Similar to results observed in the MC38 study, urine analysis indicated response to treatment several days before tumor volumes were statistically different compared to control mice (day 14 vs 17) (P ≤ 0.0001, n = 7-14, **Fig. 3i**). Collectively, these results showed that αPD1-GS indicated response to ICB treatment as early as the start of the second dose with high sensitivity and specificity.

### Protease dysregulation in tumor resistance to ICB therapy

Tumor resistance mechanisms to ICB include loss-of-function (LOF) mutations in B2M, a protein subunit of MHC-I, and JAK1, an essential signaling protein of the IFNγ response pathway^3,4^. To model resistance, we knocked out (KO) B2m or Jak1 from wildtype (WT) MC38 tumor cells with CRISPR/Cas9. We validated KO cells by TIDE (Tracking of Indels by Decomposition) analysis^33^ (**SFig. 5a**), loss of surface expression of MHC I (H2-Kb) in B2m^-/-^ cells by flow cytometry (**SFig. 5b**), reduction in GzmB and IFNγ expression by OT1 T cells after co-culture with OVA-pulsed B2m^-/-^ MC38 target cells (P ≤ 0.05, n = 3, **SFig. 5c**), and lack of upregulation of H2-Kb and PD-L1 following IFNγ stimulation of Jak1^-/-^ cells (**SFig. 5d**). To confirm resistance to ICB therapy, we treated mice bearing WT, B2m^-/-^, or Jak1^-/-^ MC38 tumors with either αPD1 or IgG1 isotype control. Whereas αPD1 treatment of WT tumors resulted in significantly smaller tumors and improved survival (MST = 30) relative to isotype control (MST = 21) (P ≤ 0.0001, n = 25, **Fig. 4a, SFig. 6**), no statistical differences in tumor burden and overall survival were observed in mice with B2m^-/-^ or Jak1^-/-^ tumors. Together, our data confirmed that LOF mutations in B2m and Jak1 render MC38 tumors resistant to αPD1 therapy.

**Figure 4.**
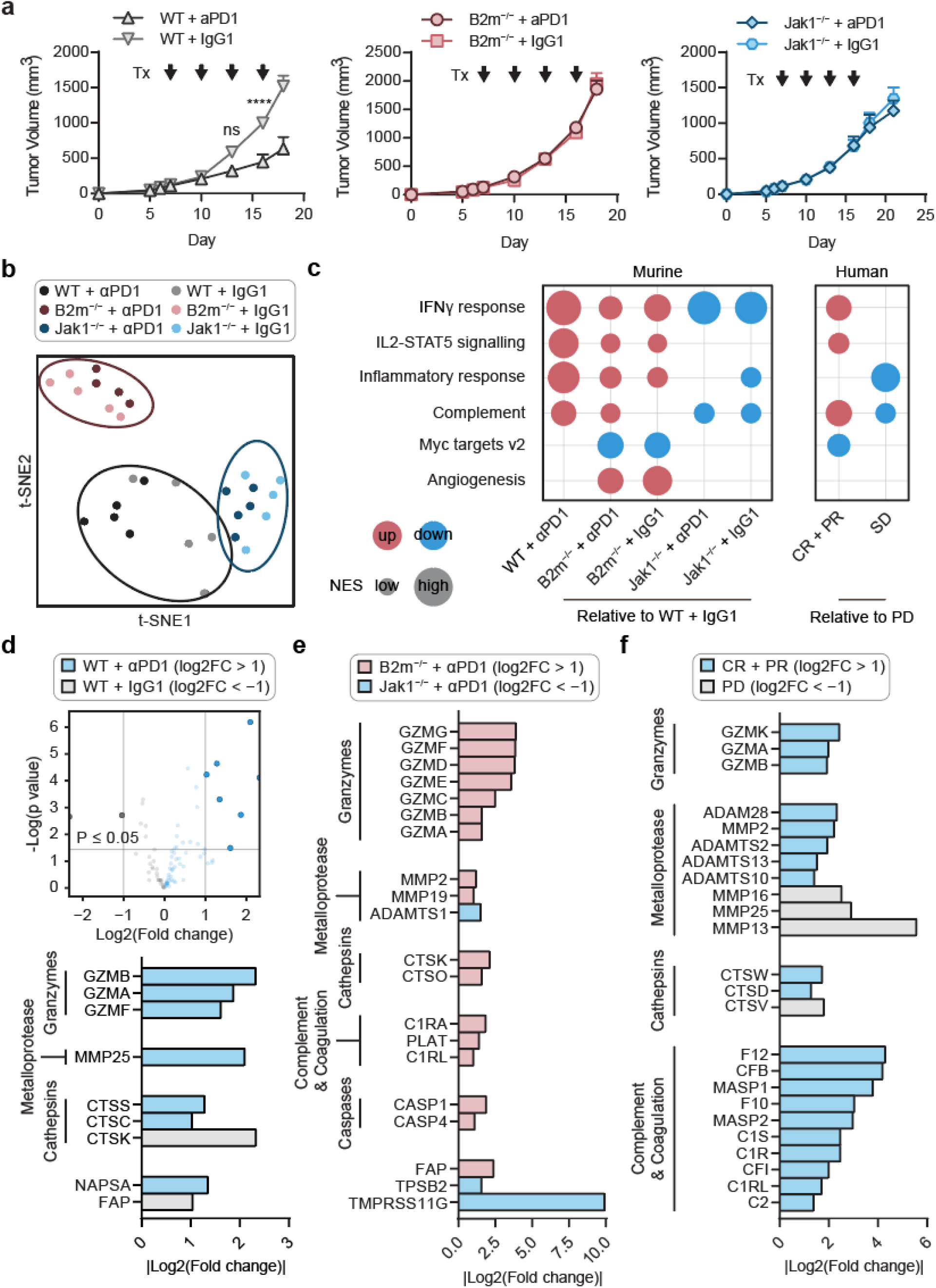
Proteases are dysregulated in ICB response and resistance. **a**, Tumor growth curves of mice bearing WT (left), B2m^-/-^ (middle), or Jak1^-/-^(right) MC38 tumor treated with αPD1 or matched IgG1 control (two-way ANOVA with Sidak’s post test and correction for multiple comparisons, ****P < 0.0001, n = 15-25). Black arrows denote the treatment time points. **b**, t-SNE plot showing global transcriptional profiles of WT, B2m^-/-^, and Jak1^-/-^ MC38 tumors treated with αPD1 or IgG1 isotype control (n = 5). **c**, Left: GSEA comparing gene set signatures of all mouse tumors and treatment groups relative to WT tumors receiving isotype control treatment (n = 5). 6 gene sets were shown from the canonical Hallmark gene sets^34^, with 4 immune- and 2 tumor-associated gene sets. Only the gene sets that are significantly different (false discovery rate < 0.05) between the two groups being compared were shown. Red color indicates upregulation in the first group, and blue indicates downregulation. The size of the circle represents the nominal enrichment score (NES). Right: similar GSEA analyses using human data from melanoma patients treated with αPD1 monotherapy^11^. Gene set signatures of the two patient groups (Complete Response (CR) + Partial Response (PR), and Stable Disease (SD)) were compared to patients with Progressive Disease (PD). **d**, Top: Volcano plots summarizing the extracellular and transmembrane proteases differentially expressed between WT MC38 tumors treated with αPD1 or IgG1 (n = 5). The threshold for differentially expressed genes (opaque dots) was defined as P value ≤ 0.05 and |log2(fold change)| ≥ 1. Bottom: waterfall plot showing the fold changes in transcript levels of proteases that are differentially expressed between these two groups. The proteases are grouped into the families of interest while the remaining are greyed out. **e**, Waterfall plot showing the fold changes in transcript levels of proteases that are differentially expressed between αPD1 treated B2m^-/-^ and Jak1^-/-^ tumors (n = 5). **f**, Waterfall plot showing the fold changes in transcript levels of proteases that are differentially expressed between human tumors from responders (CR + PR) and non-responders (PD).

To quantify the breadth of protease dysregulation in ICB response and resistance, we sequenced the transcriptomes of WT, B2m^-/-^, and Jak1^-/-^ MC38 tumors after two doses of either αPD1 or IgG1 (n = 5). By t-Distributed Stochastic Neighbor Embedding (t-SNE) analysis, we observed three distinct gene clusters corresponding to WT, B2m^-/-^, and Jak1^-/-^ tumors (**Fig. 4b**). Gene set enrichment analyses (GSEA)^34^ confirmed enrichment of immune pathways (e.g., IFNγ response, IL2-STAT5 signaling, inflammatory response, complement) in WT tumors in response to PD1 therapy, with minimal enrichment or downregulation in B2m^-/-^ and Jak1^-/-^ tumors, respectively (P ≤ 0.05, **Fig. 4c**, **SFig. 7a**). To compare with patient ICB responses, we performed GSEA on bulk tumor RNA-Seq data from advanced melanoma patients treated with αPD1 monotherapy^11^ that were classified into complete or partial responders (CR + PR), progressive disease (PD), or stable disease (SD) based on RECIST criteria^35^. We observed enrichment in immune pathways that were similar to murine tumors (e.g., IFNγ response, IL2-STAT5 signaling, complement) in CR + PR relative to PD (P ≤ 0.05, **Fig. 4c**, **SFig. 7b**).

To identify proteases dysregulated in ICB response and resistance, we compared RNA transcripts levels of WT tumors on αPD1 or IgG1 treatment and observed that the top differentially expressed proteases, as selected by a log2 fold change threshold greater than 1, were from the granzyme, metalloproteinase, and cathepsin family of enzymes (P ≤ 0.05, **Fig. 4d**, **SFig. 8a**). By comparison, B2m^-/-^ tumors on αPD1 treatment showed broader dysregulation that included proteases from the complement, coagulation, and caspase families compared to Jak1^-/-^ tumors (log2 fold change > 1, P ≤ 0.05, **Fig. 4e**, **SFig. 8b**). Similar to our mouse models, human melanoma tumors in patients^11^ that had a complete or partial response to ICB were characterized by significant upregulation of ~20 proteases across the same protease families relative to progressive disease (log2 fold change > 1, P ≤ 0.01, **Fig. 4f**). By unsupervised hierarchical clustering, protease expression profiles were primarily grouped into CR+PR compared to PD (**SFig. 8c**). Taken together, these data indicate that proteases are differentially regulated during response and resistance to ICB therapies.

### Multiplexed detection of protease activity by mass spectrometry

We next designed substrates for our INSIGHT library to detect the proteases differentially expressed in ICB response and resistance (**Fig. 5a**). We compiled published substrate sequences for five target protease families – granzymes, metalloproteases, coagulation and complement proteases, caspases, and cathepsins – and synthesized a candidate library of 66 fluorogenic substrates, which consisted of 6-11 amino acids flanked by a fluorophore (FAM) and a quencher (Dabcyl). We tested each substrate against 17 recombinant proteases (2+ per family) and quantified cleavage efficiency based on the fold change in fluorescence at 60 minutes (**Fig. 5b, SFig. 9**). To facilitate downselection, we applied t-SNE analysis and observed 4 major substrate clusters: cluster 1 contained substrates preferentially cleaved by metalloproteases, cluster 2 by metalloproteases and cathepsins, cluster 3 by coagulation and complement proteases, and cluster 4 by granzymes and caspases (**Fig. 5c**). From each cluster, we selected 3 or more representative substrates to form a final library of 14 substrates. Each substrate in this set was characterized by a 2–22 fold increase in fluorescence in the presence of target proteases (**Fig. 5d**), and the majority of substrate pairs (76%) had a Spearman’s correlation coefficient (Rs) less than 0.5, indicating low redundancy of the library (**SFig. 10**).

**Figure 5.**
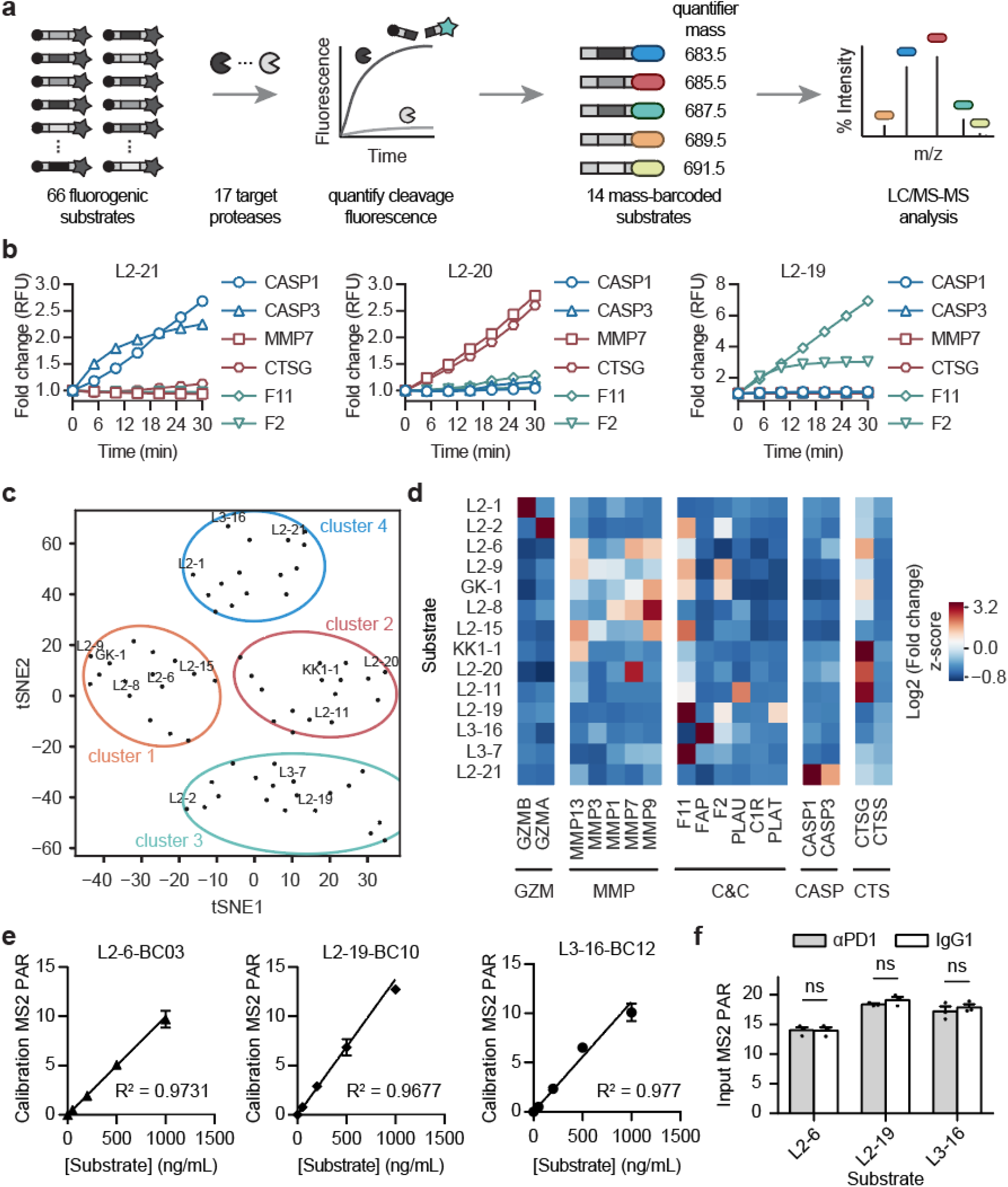
Mass-barcoded peptide sensors for multiplexed detection of protease activity. **a**, Schematic of the peptide substrate screen to identify candidate substrates for INSIGHT library. **b**, Fluorescence cleavage assays of representative substrates against recombinant proteases of interest. Each cleavage trace represents the average of 3 independent replicates. **c**, t-SNE plot showing unsupervised clustering of 66 candidate substrates into major clusters. **d**, Heat map summarizing the log2 fold change in fluorescence of 14 selected substrates at 60 min after addition of the respective recombinant protease (n = 3). Signals were row-normalized before plotting. **e**, Calibration curves of mass barcodes as quantified by LC-MS/MS. MS2 peak area from each mass barcode used to label representative substrates is normalized by peak area of an internal standard to obtain peak-area-ratio (PAR). **f**, Bar plot showing corresponding mass reporter signals (PAR) from mixtures of αPD1- or IgG1-peptide conjugates (two-way ANOVA with Tukey’s post test and correction for multiple comparisons, n = 3).

To enable multiplexed detection by mass spectrometry, we designed 14 mass barcodes by enriching the peptide reporter glutamate-fibrinopeptide B (Glufib) (EGVNDNEEGFFSAR) with different distributions of stable isotopes. As described previously^22^, this approach allows multiple reporters that share the same MS1 parent mass to be differentiated by unique quantifier MS2 fragments by tandem mass spectrometry (MS/MS) (**Table 1**). For validation, we derivatized our 14-plex substrate library with mass barcodes and confirmed that MS2 signals were linearly correlated with substrate concentrations (R^2^ ≥ 0.96, **Fig. 5e**) and that the mass barcoded substrates conjugated to αPD1 or IgG1 antibody were quantifiable after cleavage (n = 3, **Fig. 5f**). Our results showed that INSIGHT substrates are sensitive to cleavage by dysregulated proteases in the context of ICB response and resistance, and mass-barcoding allows multiplexed quantification of substrates.

**Table 1.** Mass-barcoded substrates for multiplexed urinalysis of protease activity. hF, Homophenylalanine; f, d-form phenylalanine. The barcodes are isotopically labeled Glufib peptides (EGVNDNEEGFFSAR) that share the same MS1 precursor mass for reporter pooling but produce unique fragmented MS2 quantifier ions distinguishable by liquid chromatography with tandem mass spectrometry (LC-MS/MS).

| Name | Substrate | Barcode | Precursor Mass (MS1) | Quantifier Mass (MS2) |
| --- | --- | --- | --- | --- |
| L2-1 | IEFDSG | BC01 | 806.5 | 683.5 |
| L2-2 | VANRSAS | BC02 | 808.5 | 683.5 |
| L2-6 | RPLALWRSD | BC03 | 808.5 | 685.5 |
| L2-8 | RPLGLAGK | BC04 | 808.5 | 687.5 |
| L2-9 | PLAQAVRS | BC05 | 810.5 | 683.5 |
| L2-11 | AFRFSQK | BC06 | 810.5 | 685.5 |
| L2-20 | GKPILFFRLK | BC07 | 810.5 | 687.5 |
| L2-21 | YVADAPD | BC08 | 810.5 | 689.5 |
| GK-1 | KGVPRALMVE | BC09 | 813.5 | 689.5 |
| L2-19 | fPRSGG | BC10 | 813.5 | 693.5 |
| L3-7 | EEKQRIILG | BC11 | 813.5 | 695.5 |
| L3-16 | KASGPAGPA | BC12 | 816.5 | 695.5 |
| KK1-1 | RIKFFSAQTK | BC13 | 816.5 | 697.5 |
| L2-15 | LAQA{hF}RSK | BC14 | 816.5 | 699.5 |

### Binary classification of response and resistance by 14-plex INSIGHT

To assess the potential of our 14-plex INSIGHT library to detect early on-treatment response to ICB therapy, we administered 14-plex αPD1 or IgG1 conjugates to mice bearing WT MC38 tumors at days 7, 10, and 13 (**Fig. 6a**). At each timepoint, urine samples were collected within three hours after intravenous administration and cleavage fragments were quantified by mass spectrometry. Urinary signals from dose 2 and 3 were normalized to dose 1 to account for pre-treatment baseline activity. We applied random forest classification to the data split into training and test sets by 5-fold cross validation and repeated this procedure 100 times to obtain the average area under the ROC curve (AUC)^36^. Under these conditions, INSIGHT discriminated αPD1-treated mice (n = 25) from isotype controls (n = 15) with high accuracy (AUC = 0.92 [95% CI = 0.88-0.95], sensitivity (Se) = 87%, specificity (Sp) = 86%) as early as the start of the second dose, with statistically identical classification performance at dose 3 (AUC = 0.93 [0.90-0.95], P = 0.650, paired Student’s t-test) (**Fig. 6b**). To assess the relative weight of each probe, we quantified the feature importance score and observed that probes L2-8, L3-7 and L2-1 had the largest contribution to classification accuracy with aggregate scores for dose 2 and 3 above 0.6 compared to scores of 0.3 and below for all other probes (**Fig. 6c**). These three probes were selective for granzymes, MMPs and cathepsins, including substrate L2-1 which was the same sequence previously used in αPD1-GS (**Fig. 3**). Based on the marked difference in feature importance scores, we further tested whether L2-8, L3-7, and L2-1 alone were sufficient to classify ICB responses, and found that the 3 probe set classified response with AUCs greater than 0.9 for both doses (dose 2 AUC = 0.95 [0.93-0.97]; dose 3 AUC = 0.91 [0.87-0.93]) with no statistical reduction in accuracy compared to the 14-plex panel (P = 0.147 on dose 2, P = 0.317 on dose 3, **Fig. 6d**, **SFig. 11**). These data indicated that INSIGHT discriminated ICB responders as early as the second dose with 3 probes out of the 14-plex set.

**Figure 6.**
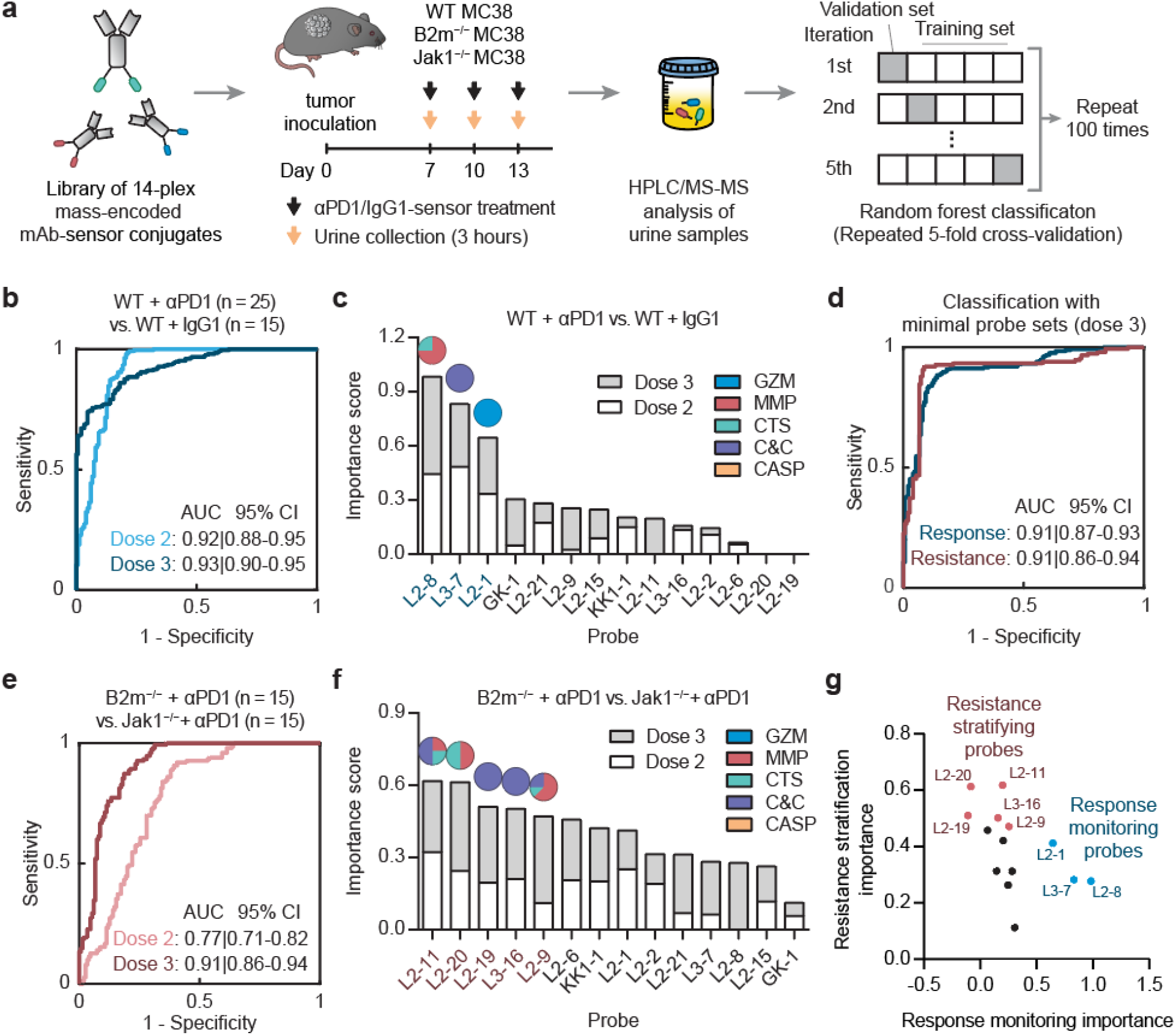
Urinary classification of ICB response and resistance. **a**, Schematic of our pipeline to develop urinary classifiers of ICB response and resistance. **b**, Area under the ROC curve (AUC) analysis showing the diagnostic specificity and sensitivity of random forest classifiers based on INSIGHT library in differentiating between αPD1-treated WT tumors (n = 25) and IgG1-treated controls (n = 15) using urine signals on dose 2 (AUC = 0.92, 95% CI = 0.88-0.95) or dose 3 (AUC = 0.93, 95% CI = 0.90-0.95). **c**, Feature importance analysis revealing the probes that are important for response monitoring. Probes with higher important scores, produced by random forest, contribute more to the diagnostic performance. The pie charts above individual probes show the protease families that are monitored by each probe. **d**, AUC analysis of random forest classifiers based on the top 3 probes (L2-8, L3-7, L2-1) for response monitoring (AUC = 0.91, 95% CI = 0.87-0.93) and the top 5 probes (L2-11, L2-20, L2-19, L3-16, and L2-9) for resistance stratification (AUC = 0.91, 95% CI = 0.86-0.94). **e**, AUC analysis of random forest classifiers based on INSIGHT library in differentiating between αPD1-treated B2m^-/-^ (n = 15) from Jak1^-/-^ MC38 (n = 15) tumors using urine signals on dose 2 (AUC = 0.77, 95% CI = 0.71-0.82) or dose 3 (AUC = 0.91, 95% CI = 0.86-0.94). **f**, Feature importance analysis revealing the probes that are important for resistance stratification. **g**, Scatter plot showing feature important scores of all 14 probes in the INSIGHT panel for response monitoring and resistance stratification. The highlighted probes belong to the minimal probe sets that achieve comparable diagnostic performance in these classification tasks as compared to using the entire INSIGHT panel.

We conducted similar longitudinal experiments to assess the ability of INSIGHT to stratify refractory tumors based on B2m^-/-^ (n = 15) or Jak1^-/-^ (n = 15) LOF mutations (**Fig. 6a**). Following urine quantification by mass spectrometry, random forest classification resulted in an AUC of 0.77 (95% CI = 0.71-0.82, Se = 84%, Sp = 65%) on dose 2, which significantly increased to 0.91 (95% CI = 0.86-0.94, Se = 87%, Sp = 81%; P ≤ 0.0001) on dose 3 (**Fig. 6e**). By feature importance analysis, we observed that a larger number of probes contributed to resistance classification where the top 5 probes had aggregate scores above 0.45 while the previous top ICB response probes, L2-8, L3-7 and L2-1, were in the bottom half by rank order (**Fig. 6f**). We further asked whether a minimal probe set could stratify resistance and by iterative analysis, we found that the top 5 probes (L2-11, L2-20, L2-19, L3-16, and L2-9) classified B2m^-/-^ from Jak1^-/-^ resistance with statistically equivalent performance to the full INSIGHT library (dose 2 AUC = 0.80 [0.74-0.84], P = 0.430; dose 3 AUC = 0.91 [0.86-0.94], P > 0.999; **Fig. 6d, SFig. 11**). Given that this subset of 5 probes did not contribute to the response monitoring classifier, we compared the importance score for all 14 probes for both classification tasks and found a strong negative correlation (R = −0.896) between the top probes for response monitoring (L2-1, L3-7, and L2-8) and stratifying resistance (L2-11, L2-20, L2-19, L3-16, and L2-9) (**Fig. 6g**). Our data indicated that binary classifiers trained on INSIGHT measurements of protease activity discriminate response and resistance to ICB therapies in mouse models.

## Discussion

In light of the central role proteases play in T cell cytotoxicity and tumor biology, our study focused on demonstrating INISIGHT as an activity-based platform to track early response and resistance to ICB therapies. We showed that αPD1-peptide conjugates act as therapeutic sensors that carry out the dual roles of reinvigorating T cell function and reporting on treatment response by the release of protease-cleaved reporters into urine for noninvasive detection. Our results with a single αPD1-GS probe to quantify GzmB activity *in vivo* showed that urinalysis of cleavage fragments anticipated response as early as the start of the second dose before tumor volumes began to diverge between treated and untreated animals. By transcriptomic analysis, we identified proteases across five families that were broadly dysregulated in tumors harboring B2m^-/-^ or Jak1^-/-^ LOF mutations. This list of proteases formed the basis of a bespoke 14-plex INSIGHT library that allowed binary classifiers trained on urine samples by machine learning to stratify the mechanism of resistance with high diagnostic accuracy. Our results support the development of activity-based biomarkers for noninvasive and longitudinal assessment of response and resistance to ICB therapies.

GzmB is the most potent pro-apoptotic granzyme and its release from granules accompanied by perforin is a primary mechanism by which CD8+ T cells exert tumoricidal activity. Compared to other tumor biomarkers (e.g., PD-L1^37^, TMB^38^, T cell-inflamed gene expression profile (GEP)^39^, microsatellite instability (MSI)^40^) and serum biomarkers (e.g., ctDNA^14,41^, TCR clonality^12,13^, memory phenotypes^12,13,42^) under investigation, GzmB is a direct biomarker of T cell cytotoxicity, and its expression has been shown to be significantly upregulated in patient tumors responsive to αPD1 and αCTLA4 therapies^43–45^. GzmB expression, therefore, has potential as an early biomarker of ICB response. Recent work on a peptide PET probe that irreversibly binds to GzmB^21,46^ demonstrated that high GzmB signals predicted early response to checkpoint therapy before changes in tumor volumes were apparent in animal models. Similarly, we observed that tumor treatment with αPD1-GS therapeutic sensors led to quantifiable levels of cleaved peptides in urine that anticipated responders from isotype controls before tumor volumes significantly diverged. As our peptide sensors are conjugated to therapeutic antibodies and administered at the time of treatment, a separate infusion of diagnostic agents is not required and response assessment can occur several hours after urine collection. In longitudinal studies with mice treated with multiple doses, we observed changes by urinalysis that indicated response as early as the start of the second dose of treatment.

GzmB expression by itself, however, is not a specific biomarker of ICB response but rather a general biomarker of T and NK cell cytotoxicity that could be elevated under confounding conditions such as reactivation of latent viruses or opportunistic infections^47–49^. Moreover, a univariate GzmB biomarker also lacks the ability to differentiate mechanisms of resistance that similarly result in loss of T cell cytotoxicity. Therefore, we investigated whether a multiplexed INSIGHT library could provide the ability to assess response and resistance to ICB therapy by multivariate classification. By transcriptomic analysis, we found that proteases are broadly dysregulated across multiple enzyme families both in tumors that respond to therapy and in tumors that harbor LOF mutations in B2M or JAK1 genes that underpin resistance to checkpoint inhibitors^3,4^. These proteases informed the design and selection of a 14-plex INSIGHT library that broadly covered protease cleavage space to provide the ability to generate high-dimensional data by mass spectrometry for classifier training. We observed that although the same INSIGHT library was used in our animal studies, separate subsets of 3 to 5 probes were ranked highest in importance depending on whether the use case was response monitoring (L2-1, L3-7, and L2-8) or stratifying resistance mechanisms (L2-11, L2-20, L2-19, L3-16, and L2-9). These probes were strongly anti-correlated (R ~ −0.9) and binary classifiers that were trained only on these minimal probe sets recapitulated the diagnostic performance of the entire 14-plex library without reductions in classification accuracy (AUROCs > 0.90). These observations lend support for a potential future strategy for human testing that involves using the same superset of probes to train separate classifiers for each intended use case. Following classifier validation, a down-selection process could then be employed to reduce the number of probes to a minimal set. This strategy may ensure the ability to generate high-dimensional data while reducing regulatory burden associated with the need to test the safety and immunogenicity of separate probe compositions.

Several key areas warrant future study. Our work outlined a general development pipeline for activity-based biomarkers that involves nomination of candidate proteases based on established biology or transcriptomic analysis, substrate design and selection, and classifier training and validation. Transcriptomic analysis of a large set of resistant tumors (i.e., primary, adaptive and acquired) with different mechanisms of action (e.g., absence of antigen presentation, insensitivity to T cells, genetic T cell exclusion^3^) would further serve to nominate differentially expressed proteases and determine the extent of conservation across cancer types and ICB therapies (e.g., αPD1 versus αCTLA-4). Given that proteases that are closely related cleave similar substrates such as the MMPs^50^, cathepsins^51^ and caspases^52^, our peptide selection process did not exclude substrates with broad selectivity for proteases within a family, which is a challenge shared by the field. This implies that assigning protease specificity to the cleavage signals will be challenging without developing probes with exquisite selectivity for target proteases, which may be possible with non-natural amino acids^53,54^, or mathematical algorithms to deconvolve complex protease signatures^55,56^. Looking forward, phase 1 studies are necessary to establish the safety of αPD1-peptide conjugates, which we anticipate to be well-tolerated in humans given its composition is similar to protease-activatable masked antibodies^29^ and T cell engagers^57^ that are undergoing clinical efficacy studies. The classifiers we described in this study are relevant for the mouse models and should not be directly mapped to humans without conducting separate training and validation studies. Overall, our results support INSIGHT as an activity-based biomarker platform to noninvasively track early response and resistance to ICB therapies from urine.

## Materials and Methods

### Animals

6-to 8-week old female mice were used at the outsets of all experiments. Pmel (B6.Cg-Thy1a/Cy Tg(TcraTcrb)8Rest/J) and OT1 (C57BL/6-Tg(TcraTcrb)1100Mjb/J) transgenic mice were bred in house using breeding pairs purchased from Jackson Lab. C57BL/6 and BALB/c mice for tumor studies were purchased from Jackson Lab. All animal procedures were approved by Georgia Tech IACUC (protocol #KWONG-A100193).

### Antibody-peptide conjugation

FITC-labelled GzmB substrate peptides ((FITC)AIEFDSGc; lower case letters = d-form amino acids) were synthesized by Tufts University Core Facility and used for *in vivo* formulations. FITC-labelled GzmB substrate peptides with internal quencher ((5-FAM)aIEFDSGK(CPQ2)kkc) were synthesized by CPC Scientific and used for all in vitro activity assays. Peptides with isobaric mass repoters were synthesized in housed using the Liberty Blue Peptide Synthesizer (CEM). Free αPD1 (kind gift of Dr. Gordon Freeman, Dana-Farber) and αCTLA4 (BioXCell; clone 9H10) antibodies were first reacted to the heterobifunctional crosslinker Succinimidyl Iodoacetate (SIA; Thermo, 5:1 molar ratio) for 2 hours at room temperature (RT) in the dark, and excess SIA were removed by buffer exchange using Amicon spin filter (30 kDa, Millipore). Cysteine-terminated peptides were mixed with mAb-SIA (10:1 molar ratio) and reacted overnight at RT in the dark to obtain mAb-peptide conjugate. The conjugates were purified on a Superdex 200 Increase 10-300 GL column using AKTA Pure FPLC System (GE Health Care). Endotoxin was removed from the samples by phase separation with Triton X-114 (Sigma) at 2% final volume ratio^58^. Final endotoxin concentrations were quantified by Pierce LAL Chromogenic Endotoxin Assay Kit (Thermo). Protein concentrations were determined by Pierce Protein Assay Kit (Thermo). Conjugates were buffered exchanged into PBS and sterile filtered before in vivo usage. Conjugation ratios of fluorescently labeled peptides were determined by corrected absorbance measurements by NanoDrop (Thermo). Conjugation of mass-encoded peptides were validated by MALDI using Autoflex mass spectrometer (Bruker).

### PD-1 binding

Binding of αPD1 conjugates to recombinant PD1 ligand was quantified using an ELISA assay developed in house, in which a high protein binding plate was coated with 1 ug/mL of recombinant Mouse PD-1 Protein (R&D, 9047-PD-100). Binding of intact αPD1-GS conjugates was quantified in a sandwich ELISA using the same PD-1 coated plate. After sample incubation, αFITC mAb (Thermo, 13-7691-82; 1:800 dilution staining concentration) was used for secondary staining. ELISA development was performed according to well-established protocol^59^.

### Circulation half-life

For half-life characterization, unconjugated αPD1 or αPD1-GS (100 ug) was administered i.v. to naïve C57BL/6 mice (Jackson Labs). At several time points following administration, blood was collected into Capillary Tubes (VWR), and serum was isolated by centrifugation. Serum concentrations of unconjugated αPD1 and αPD1-GS were determined by the PD1 binding and intact PD1 ELISA respectively.

### Recombinant protease cleavage assays

αPD1 was conjugated with GzmB peptide substrates carrying an internal CPQ2 quencher to allow cleavage detection by fluorescent measurements. αPD1-GS (1.3 uM by peptide) was incubated in PBS at 37 °C with fresh mouse serum, murine Granzyme B (0.17 μM; Peprotech), human thrombin (13.5 μM; HaemTech), mouse thrombin (12.5 μM; HaemTech), cathepsin B (1.5 μM, R&D), C1r (1.43 μM; Sigma), C1s (1.80 μM; Sigma), MMP9 (0.1 μM, R&D). Sample fluorescence was measured for 60 minutes using Cytation 5 plate reader (Biotek).

### Sensing protease activity during T cell killing

B16-F10 cells (ATCC) were cultured in DMEM supplemented with 10% FBS and 1% penicillin-streptomycin (Thermo). CD8+ T cells were isolated from either OT1 or Pmel (Jackson Labs) splenocytes by MACS using CD8a Microbeads (Miltenyi). Cells were activated by seeding in 96-well plates pre-coated with anti-mouse CD3e (1 μg/ml working concentration, Clone: 145-2C11, BD) and anti-mouse CD28 (2 μg/ml working concentration, Clone: 37.51, BD) at 2×10^6^ cells/ml in RPMI 1640 supplemented with 10% FBS, 100U/ml penicillin-streptomycin, 1X non-essential amino acids (Gibco), 1mM sodium pyruvate, 0.05mM 2-mercaptoethanol, and 30U/ml hIL-2 (Roche). After 2 days, cells were washed and transferred to untreated culture flasks for expansion. Between day 4 to 6 after activation, activated T cells were washed before coincubated with 3×10^4^ B16 target cells at various T cell to effector cell ratios. After 48 hours, coculture supernatants were collected for LDH and GzmB measurements by the Pierce LDH Cytotoxicity Assay Kit (Thermo) and GzmB Mouse ELISA Kit (Thermo, BMS6029) respectively. To assess sensor activation during T cell killing, cocultured of T cells and target cells were spiked in with either αPD1-GS, αPD1 conjugated with control peptide (LQRIYK), and unconjugated αPD1. After 48 hours, fluorescence of coculture supernatant were measured using Cytation 5 plate reader (Biotek).

### Tumor models

CT26 (ATCC), MC38 (kind gift of the NCI and Dr. Dario Vignali, University of Pittsburgh), and B2m^-/-^ vs. Jak1^-/-^ MC38 tumor cells were cultured in DMEM supplemented with 10% FBS and 1% penicillin-streptomycin (Thermo). Cells were grown to a good density (~70% confluence) before trypsinized for tumor inoculation. On the day of inoculation, C57BL/6 and BALB/c mice were shaved and injected s.c. into the left flank with either 1×10^6^ MC38 or CT26 cells respectively. Tumor burden were monitored until average tumor volume, quantified as 0.52 x length x width x depth, was approximately 100 mm^3^ before initiating treatment. Mice were administered with αPD1 and/or αCTLA4 antibody-sensor conjugates or matched isotype control (100-150 ug/injection) every 3 or 4 days.

### Flow cytometry analysis of intratumoral T cells

Tumor dissociation and staining for flow cytometry. Less than 1g of murine tumors were enzymatically and mechanically dissociated using Mouse Tumor Dissociation Kit (Miltenyi) and gentleMACS Dissociator (Miltenyi). TILs were then isolated from the single cell suspension using a density gradient with Percoll Centrifugation Media (GE Life Sciences) and DMEM Media (10% FBS, 1% Penstrep) at 44:56 volume ratio. TILs were counted with Trypan Blue (Thermo), and approximately 1×10^6^ viable cells per sample were stained for flow cytometry analysis. Cells were first stained for surface markers in FACS Buffer (1x DPBS, 2% FBS, 1 mM EDTA, 25 mM HEPES). Intracellular staining was performed using eBioscience Intracellular Fixation & Permeabilization Buffer Set (Thermo). All antibodies were used for staining at 1:100 dilution from stock concentrations. Stained cells were analyzed by LSRFortessa Flow Cytometer (BD).

Antibody clones. CD45 (30-F11), CD8 (53-6.7), CD44 (IM7), PD-1 (29F.1A12), TIM3 (RMT3-23), CD4 (RM4-5), NK1.1 (PK136), CD19 (6D5), GZMB (GB12). Viability was accessed by staining with LIVE/DEAD Fixable Dye (Thermo).

### Urinary detection of therapeutic response and resistance to ICB therapy

At 3 hours after administration of ICB antibody-sensor conjugates, urine was collected and analyzed for noninvasive detection of therapeutic response and resistance. FITC reporters were isolated from urine samples using Dynabeads (Thermo) decorated with αFITC antibody (Genetex). Sample fluorescence was measured by Cytation 5 plate reader (Biotek), and reporter concentrations were determined by using a known FITC ladder. Concentrations of isobaric mass reporters were quantified by Syneous Health (Morrisville, NC) using LC-MS/MS.

### Cas9 knockout of B2m and Jak1

CRISPR guide RNA’s were designed to target two exons in either B2m (g1: GACAAGCACCAGAAAGACCA, g2: GGATTTCAATGTGAGGCGGG) or Jak1 (g1: GTGAACTGGCATCAAGGAGT, g2: GCTTGGTGCTCTCATCGTAC) in the Mus musculus GRCm38 genome. Top and bottom guide oligonucleotides were annealed using T4 PNK (NEB) and ligated into the backbone of eSpCas9_PuroR_GFP plasmid (Sigma) using BbsI cut sites and T7 ligase (NEB). 1×10^5^ MC38 cells were transfected with gRNA-ligated eSpCas9 plasmids for 48 hours using TransIT-LT1 transfection reagent (Mirus Bio) in Opti-MEM (Thermo Fisher) and cultured for 3 passages in DMEM supplemented with 10% FBS and 1% penicillin-streptomycin (D10). Selection of transfected cells were done by supplementing culture media with 2 ug/mL puromycin (Thermo Fisher). Cells incubated with B2m-directed guides were stained with anti-mouse H-2Kb (clone AF6-88.5). H-2Kb-negative GFP-positive cells were sorted into single cells on a 96-well plate using FACSAria Fusion (BD Biosciences) and cultured for 2-3 weeks in D10. For cells incubated with Jak1-directed guides, GFP-positive cells were sorted into single cells and cultured for 2-3 weeks in D10. Clones that passed the functional assays for successful deletion of B2m or Jak1 are selected for tumor studies.

### In vitro validation

DNA was isolated from single-cell WT and knockout clones, and a PCR reaction was done to amplify the edited regions within B2m and Jak1 exons. The PCR products were sequenced by Sanger sequencing, and sequencing results were analyzed with TIDE (Tracking of Indels by Decomposition) analysis^33^ to confirm knockout efficiency. WT and knockout tumor cells were stained for H2-Kb (clone AF6-88.5) to confirm the functional loss of B2m. WT and B2m^-/-^ were pulsed with SIINFEKL (30 uM peptide concentration), washed, and coincubated with plate-activated OT1 T cells at 5:1 ratio of effector:target cell. After overnight incubation, cells were washed and stained for CD8 (53-6.7), IFNγ (XMG1.2), and GzmB (GB12). For IFNγ stimulation assay, WT and knockout tumor cells were incubated with recombinant murine IFNγ (Peprotech; 500 EU/mL) for 2 days and stained for surface expression of H2-Kb (AF6-88.5) and PD-L1 (10F.9G2).

### Tumor RNA isolation and sequencing

Mice bearing WT, B2m^-/-^, Jak1^-/-^ MC38 tumors were treated with either αPD1 or IgG1 (100 ug) every 3 or 4 days. After the third administration, approximately 50 mg of tumors were dissected and rapidly frozen with dry ice and IPA. Frozen tumor samples were homogenized in MACS M Tubes (Miltenyi) using the MACS Dissociator (Miltenyi). Total RNA was isolated from the homogenate using the RNeasy Plus Mini Kit (Qiagen). Library preparation with TruSeq RNA Library Prep Kit (Illumina) and mRNA NGS sequencing (40×10^6^ paired end read) were performed by Admera Health (South Plainfield, NJ).

### RNA-seq data mapping and visualization

Raw FASTQ reads passing quality control (FastQC v0.11.2) were aligned on the mm10 reference genome using STAR aligner (v2.5.2a) with default parameters. Aligned fragments were then counted and annotated using Rsamtools (v3.2) and Cufflinks (v.2.2.1) after a ‘dedup’ step using BamUtils (v1.0.11). t-SNE embedding results were performed in sklearn (v0.23.1) using all murine genes. Heat maps were plotted with seaborn’s (v.0.9.0) clustermap function. Rows were gaussian normalized, and the dendrograms shown for clustering come from hierarchical clustering using Euclidean distance as a metric.

### Differential expression and gene set enrichment analysis

Differential expression was performed using the edgeR package (v3.24.3) in R using the exactTest method with tagwise dispersion. For mouse data, TMM normalization considering mice in all treatment groups was performed to remove library size effect through the calcNormFactors function. For human data^11^, TMM normalization was performed using the two groups being compared. For both datasets, differential expression was performed on Ensembl IDs before mapping to gene names. Then the identified differentially expressed genes were filtered by a list of extracellular and transmembrane endopeptidases queried from UniProt. Gene set enrichment analysis (GSEA) was performed using the fgsea package (v1.8.0) in R. To rank genes, differential expression analysis was first performed on the entire gene set. Genes are then ranked by −sign(logFC)*log(pval). Hallmark gene sets (MSigDB) were used for all GSEA analyses.

### Peptide substrate synthesis

To optimize peptide substrates for target proteases, a library of potential substrates flanked by 5FAM fluorescent dye and DABCYL quencher (5FAM-substrate-Lys{DABCYL}-Amide) was synthesized by Genscript or manufactured in-house using Liberty Blue peptide synthesizer (CEM). The peptide synthesis scale used was 0.025 mM, and Low-loading rink amide resin (CEM) was used. Amino acids (Chem-Impex) were resuspended in DMF (0.08 M), as were all synthesis buffers. Activator buffer used was Diisopropylcarbodiimide (DIC; Sigma) (0.25 M) and the activator base buffer was Oxyma (0.25 M; CEM) while the deprotection buffer was Piperidine (20%; Sigma) supplemented with Oxyma (0.1 M). Crude peptides were purified on 1260 Infinity II HPLC system (Agilent) until a purity of 80% was achieved. Peptide mass and purity were validated by LC-MS (Agilent) and Autoflex TOF mass spectrometer (Bruker).

### Protease substrate library optimization

Fluorescently quenched peptide substrates (10 uM) were incubated in manufacturer-recommended buffers at 37°C with recombinant proteases (25 nM). Our set of human recombinant proteases included Granzyme A, Granzyme B, MMP1, MMP3, MMP7, MMP9, MMP13, Caspase 1, Caspase 3, Cathepsin G, Cathepsin S (Enzo), human thrombin, human Factor XIa (HaemTech), C1R, Fibroblast Activation Protein alpha/FAP, t-Plasminogen Activator/tPA Protein, and u-Plasminogen Activator/Urokinase (R&D systems). Sample fluorescence (Ex/Em = 488 nm/525 nm) were measured for 180 minutes using Cytation 5 plate reader (Biotek). Enzyme cleavage rates were quantified as relative fluorescence increase over time normalized to fluorescence before addition of protease. Hierarchical clustering was performed in python, using log2 fluorescence fold change at 60 minutes. A positive cleavage event was defined as having fluorescence signal more than 2-fold above background. Correlation analysis with Spearman coefficient was done on the cleavage patterns of all peptide substrates for selection of 14 substrates for library construction. These peptide substrates were paired with isobaric mass reporters based on the GluFib peptide (Table 1) and synthesized using Liberty Blue peptide synthesizer (CEM).

### Urinary differentiation of ICB resistant mechanisms

Random forest was used to train classifiers based on urinary reporter signals that differentiate therapeutic response and stratify resistant mechanisms. Response monitoring classifiers were trained on reporter concentration whereas resistance stratifying classifiers were trained on mean normalized reporter concentration. All urine signals were normalized on a per mouse basis by signals on the first dose to performed paired sample analyses. For each classification task, we used five-fold cross validation by randomly left out 1/5^th^ samples as the test set and used the remaining samples as training sets. This process was repeated 100 times, and the final performance was generated as the average area under the ROC curve (AUROC) for all train-test results. Comparisons between diagnostic performance was done by two-way paired t-test.

### Software and Statistical Analysis

Graphs were plotted and appropriate statistical analyses were conducted using GraphPad Prism (*P< 0.05, **P < 0.01, ***P < 0.001, ****P < 0.0001; central values depict the means, and error bars depict s.e.m.). Measurements were taken from distinct samples. Flow cytometry data were analyzed using FlowJo X (FlowJo, LLC). Power analyses were performed using G*Power 3.1 (HHUD).

## Data availability

All data supporting the findings of this study are available in the manuscript and its Supplementary Information. Requests for raw data can be addressed to the corresponding author.

## Code availability

All codes used in the manuscript are available upon request to the corresponding author.

## Acknowledgments

This work was funded by the NIH Director’s New Innovator Award DP2HD091793 and the National Cancer Institute R01 grant 5R01CA237210. Q.D.M. and A.S. are supported by the NSF Graduate Research Fellowships Program (Grant No. DGE-1650044). G.A.K. holds a Career Award at the Scientific Interface from the Burroughs Wellcome Fund. P.Q. is an ISAC Marylou Ingram Scholar and a Carol Ann and David D. Flanagan Faculty Fellow. This work was performed in part at the Georgia Tech Institute for Electronics and Nanotechnology, a member of the National Nanotechnology Coordinated Infrastructure, which is supported by the National Science Foundation (Grant ECCS-1542174). This content is solely the responsibility of the authors and does not necessarily represent the official views of the National Institutes of Health. The authors would like to thank the staffs at Georgia Tech mass spectrometry core, flow cytometry analysis core, and the animal facility for their assistance in performing our studies.

## Author contributions

Q.D.M., J.R.B, and G.A.K. conceived of the idea. Q.D.M., C.X., J.R.B, A.S., H.P., F-Y.S., S.Z.S., P.Q., and G.A.K. designed experiments and interpreted results. Q.D.M., C.X., J.R.B, A.S., H.P., F-Y.S., S.Z.S., H.S., A.M.H., and T.T.L. carried out the experiments. Q.D.M., A.S., and G.A.K. wrote the manuscript.

**Correspondence and requests for materials** should be addressed to the corresponding author (G.A.K).

### Ethical compliance

All authors have complied with relevant ethical regulations while conducting this study.

## References

1. Ribas, A. & Wolchok, J. D. Cancer immunotherapy using checkpoint blockade. Science 359, 1350–1355 (2018).

2. Sharma, P. & Allison, J. P. The future of immune checkpoint therapy. Science 348, 56–61 (2015).

3. Sharma, P., Hu-Lieskovan, S., Wargo, J. A. & Ribas, A. Primary, Adaptive, and Acquired Resistance to Cancer Immunotherapy. Cell 168, 707–723 (2017).

4. Kalbasi, A. & Ribas, A. Tumour-intrinsic resistance to immune checkpoint blockade. Nat. Rev. Immunol. 1–15 (2019) doi:10.1038/s41577-019-0218-4.

5. Nishino, M., Ramaiya, N. H., Hatabu, H. & Hodi, F. S. Monitoring immune-checkpoint blockade: response evaluation and biomarker development. Nat. Rev. Clin. Oncol. 14, 655–668 (2017).

6. Hodi, F. S. et al. Evaluation of Immune-Related Response Criteria and RECIST v1.1 in Patients With Advanced Melanoma Treated With Pembrolizumab. J. Clin. Oncol. 34, 1510–1517 (2016).

7. Garon, E. B. et al. Pembrolizumab for the Treatment of Non–Small-Cell Lung Cancer. N. Engl. J. Med. 372, 2018–2028 (2015).

8. Nishino, M. et al. Immune-Related Tumor Response Dynamics in Melanoma Patients Treated with Pembrolizumab: Identifying Markers for Clinical Outcome and Treatment Decisions. Clin. Cancer Res. 23, 4671–4679 (2017).

9. Gerwing, M. et al. The beginning of the end for conventional RECIST — novel therapies require novel imaging approaches. Nat. Rev. Clin. Oncol. 16, 442–458 (2019).

10. Mandal, R. & Chan, T. A. Personalized Oncology Meets Immunology: The Path toward Precision Immunotherapy. Cancer Discov. 6, 703–713 (2016).

11. Riaz, N. et al. Tumor and Microenvironment Evolution during Immunotherapy with Nivolumab. Cell 171, 934–949.e16 (2017).

12. Fairfax, B. P. et al. Peripheral CD8 + T cell characteristics associated with durable responses to immune checkpoint blockade in patients with metastatic melanoma. Nat. Med. 26, 193–199 (2020).

13. Valpione, S. et al. Immune awakening revealed by peripheral T cell dynamics after one cycle of immunotherapy. Nat. Cancer 1, 210–221 (2020).

14. Goldberg, S. B. et al. Early Assessment of Lung Cancer Immunotherapy Response via Circulating Tumor DNA. Clin. Cancer Res. 24, 1872–1880 (2018).

15. Kessenbrock, K., Plaks, V. & Werb, Z. Matrix Metalloproteinases: Regulators of the Tumor Microenvironment. Cell 141, 52–67 (2010).

16. Dudani, J. S., Warren, A. D. & Bhatia, S. N. Harnessing Protease Activity to Improve Cancer Care. Annu. Rev. Cancer Biol. 2, 353–376 (2018).

17. Martínez-Lostao, L., Anel, A. & Pardo, J. How Do Cytotoxic Lymphocytes Kill Cancer Cells? Clin. Cancer Res. 21, 5047–5056 (2015).

18. Hilderbrand, S. A. & Weissleder, R. Near-infrared fluorescence: application to in vivo molecular imaging. Curr. Opin. Chem. Biol. 14, 71–79 (2010).

19. Sanman, L. E. & Bogyo, M. Activity-Based Profiling of Proteases. Annu. Rev. Biochem. 83, 249–273 (2014).

20. Savariar, E. N. et al. Real-time In Vivo Molecular Detection of Primary Tumors and Metastases with Ratiometric Activatable Cell-Penetrating Peptides. Cancer Res. 73, 855–864 (2013).

21. Larimer, B. M. et al. Granzyme B PET Imaging as a Predictive Biomarker of Immunotherapy Response. Cancer Res. 77, 2318–2327 (2017).

22. Kwong, G. A. et al. Mass-encoded synthetic biomarkers for multiplexed urinary monitoring of disease. Nat. Biotechnol. 31, 63–70 (2013).

23. Lin, K. Y., Kwong, G. A., Warren, A. D., Wood, D. K. & Bhatia, S. N. Nanoparticles That Sense Thrombin Activity As Synthetic Urinary Biomarkers of Thrombosis. ACS Nano 7, 9001–9009 (2013).

24. Warren, A. D., Kwong, G. A., Wood, D. K., Lin, K. Y. & Bhatia, S. N. Point-of-care diagnostics for noncommunicable diseases using synthetic urinary biomarkers and paper microfluidics. Proc. Natl. Acad. Sci. 111, 3671–3676 (2014).

25. Kwong, G. A. et al. Mathematical framework for activity-based cancer biomarkers. Proc. Natl. Acad. Sci. 112, 12627–12632 (2015).

26. Mac, Q. D. et al. Non-invasive early detection of acute transplant rejection via nanosensors of granzyme B activity. Nat. Biomed. Eng. 3, 281–291 (2019).

27. Kirkpatrick, J. D. et al. Urinary detection of lung cancer in mice via noninvasive pulmonary protease profiling. Sci. Transl. Med. 12, (2020).

28. Efremova, M. et al. Targeting immune checkpoints potentiates immunoediting and changes the dynamics of tumor evolution. Nat. Commun. 9, 32 (2018).

29. Desnoyers, L. R. et al. Tumor-Specific Activation of an EGFR-Targeting Probody Enhances Therapeutic Index. Sci. Transl. Med. 5, 207ra144–207ra144 (2013).

30. Strohl, W. R. Fusion Proteins for Half-Life Extension of Biologics as a Strategy to Make Biobetters. BioDrugs 29, 215–239 (2015).

31. Duraiswamy, J., Kaluza, K. M., Freeman, G. J. & Coukos, G. Dual Blockade of PD-1 and CTLA-4 Combined with Tumor Vaccine Effectively Restores T-Cell Rejection Function in Tumors. Cancer Res. 73, 3591–3603 (2013).

32. Selby, M. J. et al. Preclinical Development of Ipilimumab and Nivolumab Combination Immunotherapy: Mouse Tumor Models, In Vitro Functional Studies, and Cynomolgus Macaque Toxicology. PLOS ONE 11, e0161779 (2016).

33. Brinkman, E. K., Chen, T., Amendola, M. & van Steensel, B. Easy quantitative assessment of genome editing by sequence trace decomposition. Nucleic Acids Res. 42, e168–e168 (2014).

34. Liberzon, A. et al. The Molecular Signatures Database Hallmark Gene Set Collection. Cell Syst. 1, 417–425 (2015).

35. Schwartz, L. H. et al. RECIST 1.1 – Update and Clarification: From the RECIST Committee. Eur. J. Cancer Oxf. Engl. 1990 62, 132–137 (2016).

36. Arlot, S. & Celisse, A. A survey of cross-validation procedures for model selection. Stat. Surv. 4, 40–79 (2010).

37. Patel, S. P. & Kurzrock, R. PD-L1 Expression as a Predictive Biomarker in Cancer Immunotherapy. Mol. Cancer Ther. 14, 847–856 (2015).

38. Chan, T. A. et al. Development of tumor mutation burden as an immunotherapy biomarker: utility for the oncology clinic. Ann. Oncol. 30, 44–56 (2019).

39. Cristescu, R. et al. Pan-tumor genomic biomarkers for PD-1 checkpoint blockade–based immunotherapy. Science 362, (2018).

40. Chang, L., Chang, M., Chang, H. M. & Chang, F. Microsatellite Instability: A Predictive Biomarker for Cancer Immunotherapy. Appl. Immunohistochem. Mol. Morphol. 26, e15 (2018).

41. Bratman, S. V. et al. Personalized circulating tumor DNA analysis as a predictive biomarker in solid tumor patients treated with pembrolizumab. Nat. Cancer 1, 873–881 (2020).

42. Tietze, J. K. et al. The proportion of circulating CD45RO+CD8+ memory T cells is correlated with clinical response in melanoma patients treated with ipilimumab. Eur. J. Cancer 75, 268–279 (2017).

43. Chen, P.-L. et al. Analysis of Immune Signatures in Longitudinal Tumor Samples Yields Insight into Biomarkers of Response and Mechanisms of Resistance to Immune Checkpoint Blockade. Cancer Discov. 6, 827–837 (2016).

44. Tumeh, P. C. et al. PD-1 blockade induces responses by inhibiting adaptive immune resistance. Nature 515, 568–571 (2014).

45. Jiang, P. et al. Signatures of T cell dysfunction and exclusion predict cancer immunotherapy response. Nat. Med. 24, 1550–1558 (2018).

46. Larimer, B. M. et al. The Effectiveness of Checkpoint Inhibitor Combinations and Administration Timing Can Be Measured by Granzyme B PET Imaging. Clin. Cancer Res. 25, 1196–1205 (2019).

47. Zhang, X. et al. Hepatitis B virus reactivation in cancer patients with positive Hepatitis B surface antigen undergoing PD-1 inhibition. J. Immunother. Cancer 7, 322 (2019).

48. Del Castillo, M. et al. The Spectrum of Serious Infections Among Patients Receiving Immune Checkpoint Blockade for the Treatment of Melanoma. Clin. Infect. Dis. 63, 1490–1493 (2016).

49. Fujita, K. et al. Emerging concerns of infectious diseases in lung cancer patients receiving immune checkpoint inhibitor therapy. Respir. Med. 146, 66–70 (2019).

50. Aguilera, T. A., Olson, E. S., Timmers, M. M., Jiang, T. & Tsien, R. Y. Systemic *in vivo* distribution of activatable cell penetrating peptides is superior to that of cell penetrating peptides. Integr. Biol. 1, 371–381 (2009).

51. Whitley, M. J. et al. A mouse-human phase 1 co-clinical trial of a protease-activated fluorescent probe for imaging cancer. Sci. Transl. Med. 8, 320ra4–320ra4 (2016).

52. Timmer, J. C. & Salvesen, G. S. Caspase substrates. Cell Death Differ. 14, 66–72 (2007).

53. Poreba, M. et al. Unnatural amino acids increase sensitivity and provide for the design of highly selective caspase substrates. Cell Death Differ. 21, 1482–1492 (2014).

54. Rut, W. et al. Recent advances and concepts in substrate specificity determination of proteases using tailored libraries of fluorogenic substrates with unnatural amino acids. Biol. Chem. 396, 329–337 (2015).

55. Miller, M. A. et al. Proteolytic Activity Matrix Analysis (PrAMA) for simultaneous determination of multiple protease activities. Integr. Biol. 3, 422–438 (2011).

56. Zhuang, Q., Holt, B. A., Kwong, G. A. & Qiu, P. Deconvolving multiplexed protease signatures with substrate reduction and activity clustering. PLOS Comput. Biol. 15, e1006909 (2019).

57. Austin, R. J. et al. TriTACs, a Novel Class of T-Cell–Engaging Protein Constructs Designed for the Treatment of Solid Tumors. Mol. Cancer Ther. 20, 109–120 (2021).

58. Triplett, T. A. et al. Reversal of indoleamine 2,3–dioxygenase-mediated cancer immune suppression by systemic kynurenine depletion with a therapeutic enzyme. Nat. Biotechnol. 36, 758–764 (2018).

59. Clark, M. F., Lister, R. M. & Bar-Joseph, M. ELISA techniques. in Methods in Enzymology vol. 118 742–766 (Academic Press, 1986).

