## Supplemental Materials for "Activity-based urinary biomarkers of response and resistance to checkpoint blockade immunotherapy"

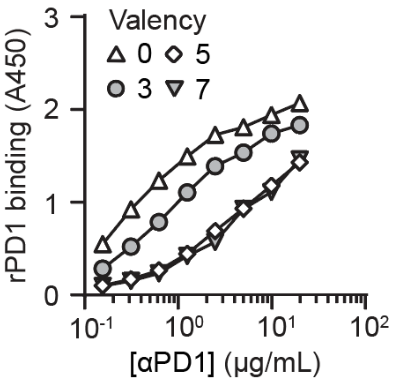


**Supplementary Figure S1 | The effect of peptide valency to antibody binding.** ELISA assays comparing the binding affinity of αPD1-GS with different peptide to antibody ratios to unmodified αPD1 antibody (unfilled uppointing triangle).

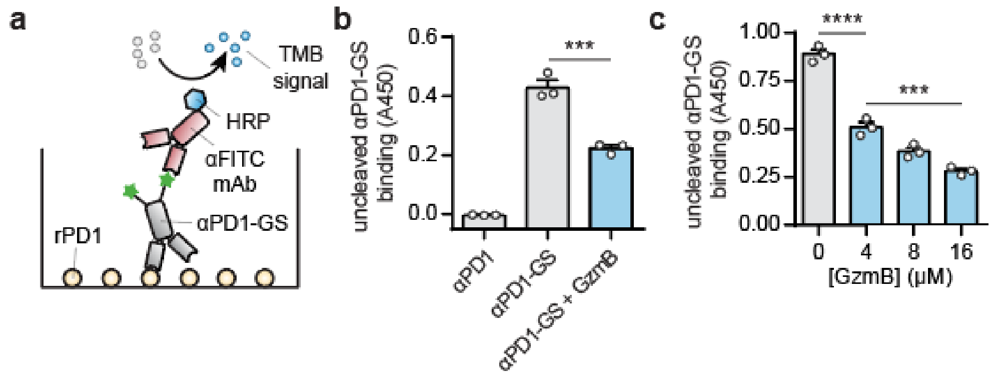


**Supplemental Figure S2 | Characterization of uncleaved αPD1-GS by sandwich ELISA assay. a**, Schematics of the assay, which requires the binding of αPD1 antibody-peptide conjugate to plate-coated recombinant PD1 protein (rPD1) and the binding of αFITC secondary antibody to uncleaved FITC-labeled peptides on the conjugate to emit a detection signal. **b**, ELISA assays showing detection signals (absorbance at 450 nm) of αPD1, αPD1-GS, and αPD1-GS in presence of recombinant GzmB (one-way ANOVA with Dunnett's post test and correction for multiple comparison, ***$P < 0.001$, $n = 3$). **c**, ELISA assays showing detection signals of αPD1-GS in presence of no or various concentrations of recombinant GzmB (one-way ANOVA with Turkey's post test and correction for multiple comparison, ****$P < 0.0001$, $n = 3$).

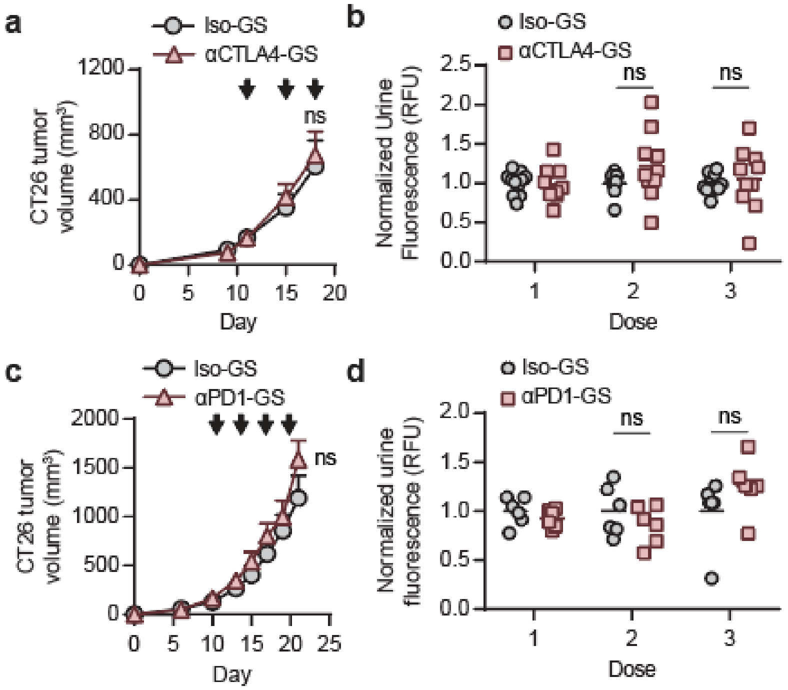


**Supplemental Figure S3 | Diagnostic performance of αPD1-GS in ICB nonresponsive models. a,** Tumor growth curves of CT26 tumor bearing mice treated with either αCTLA4-GS or matched IgG2 isotype control (Iso-GS) (two-way ANOVA with Sidak's post test and correction for multiple comparisons, ns = not significant, n = 10-11). Black arrows denote the treatment time points. **b**, Normalized urine fluorescence of mice with CT26 tumors after each administration of αCTLA4-GS or Iso-GS (two-way ANOVA with Sidak's post test and correction for multiple comparisons, ns = not significant, n = 10-11). **c,** Tumor growth curves of CT26 tumor bearing mice treated with αPD1-GS or matched IgG1 isotype control (Iso-GS) (two-way ANOVA with Sidak's post test and correction for multiple comparisons, ns = not significant, n = 6). Black arrows denote the treatment time points. **d**, Normalized urine fluorescence of mice with CT26 tumors after each administration of αPD1-GS or Iso-GS (two-way ANOVA with Sidak's post test and correction for multiple comparisons, ns = not significant, n = 6).

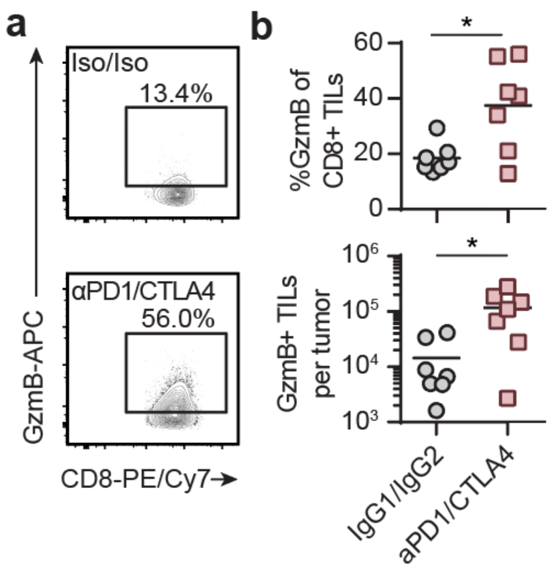


**Supplemental Figure S4 | Flow cytometry analysis of tumor infiltrating lymphocytes from CT26 tumors treated with ICB combination therapy. a**, Flow cytometry plots showing intracellular GzmB expression of CD8+ TILs from CT26 tumors treated with αPD1-GS and αCTLA4 combination therapy or matched isotype control conjugated with the GzmB peptide substrates (Iso-GS/Iso). **b**, Quantified plots showing percentages of GzmB+ cells within the CD8+ TILs or the numbers of GzmB+CD8+ TILs that were isolated from CT26 tumors treated with combination therapy or matched isotype control (two-sided Student's t-test, *$P < 0.05$, $n = 7$).

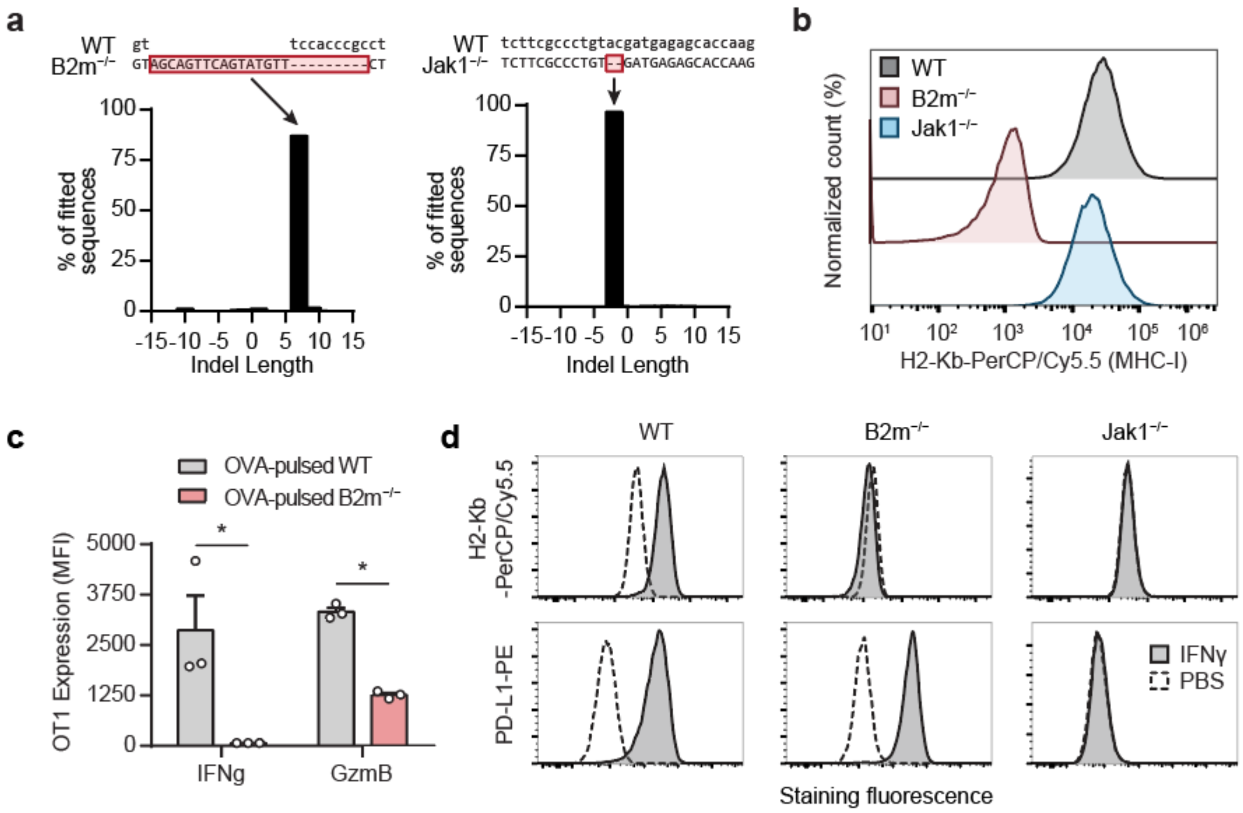


**Supplemental Figure S5 | In vitro characterization of B2m$^{-/-}$ and Jak1$^{-/-}$ MC38 tumor cells.** **a**, Sequencing alignment and TIDE (Tracking of Indels by Decomposition) analyses of MC38 tumor cells after CRISPR/Cas9 editing of B2m (left) or Jak1 (right). **b**, Flow cytometry histograms showing the staining of H2-Kb on WT, B2m$^{-/-}$, and Jak1$^{-/-}$ MC38 tumor cells. **c**, Bar plots showing median fluorescence intensity (MFI) of T cell effector molecules IFNγ and GzmB expressed by OT1 transgenic T cells in cocultures with wildtype (WT) or B2m$^{-/-}$ MC38 tumor cells pulsed with the cognate antigen ovalbumin (OVA) (two-tailed Student's t-test, n = 3). **d**, Flow cytometry histograms showing expression of MHC-I (H2-Kb) and PD-L1 on the surface of WT, B2m$^{-/-}$, and Jak1$^{-/-}$ MC38 tumor cells upon stimulation with either IFNγ or PBS control.

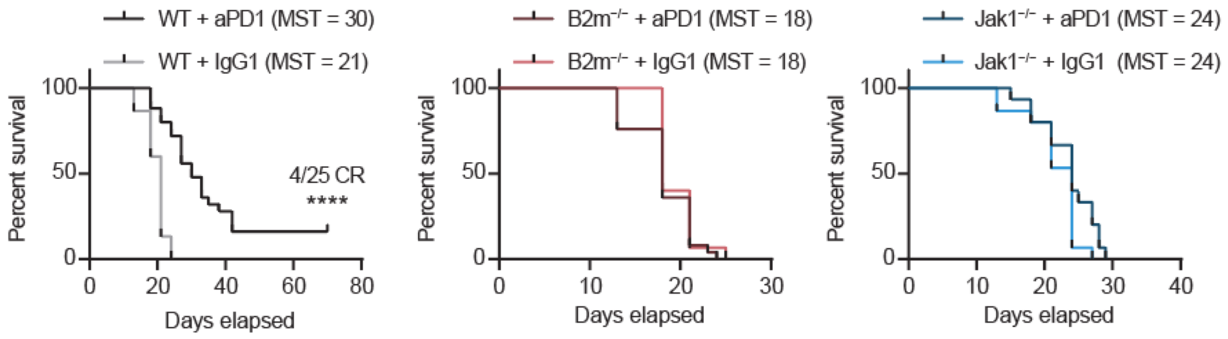


**Supplemental Figure S6 | Survival analysis of WT and knockout tumors treated with αPD1 monotherapy.** Survival curves of mice bearing WT (left), B2m$^{-/-}$ (middle), or Jak1$^{-/-}$(right) MC38 tumor treated with αPD1 or matched isotype control (Log-rank (Mantel-Cox) test, n = 15-25).

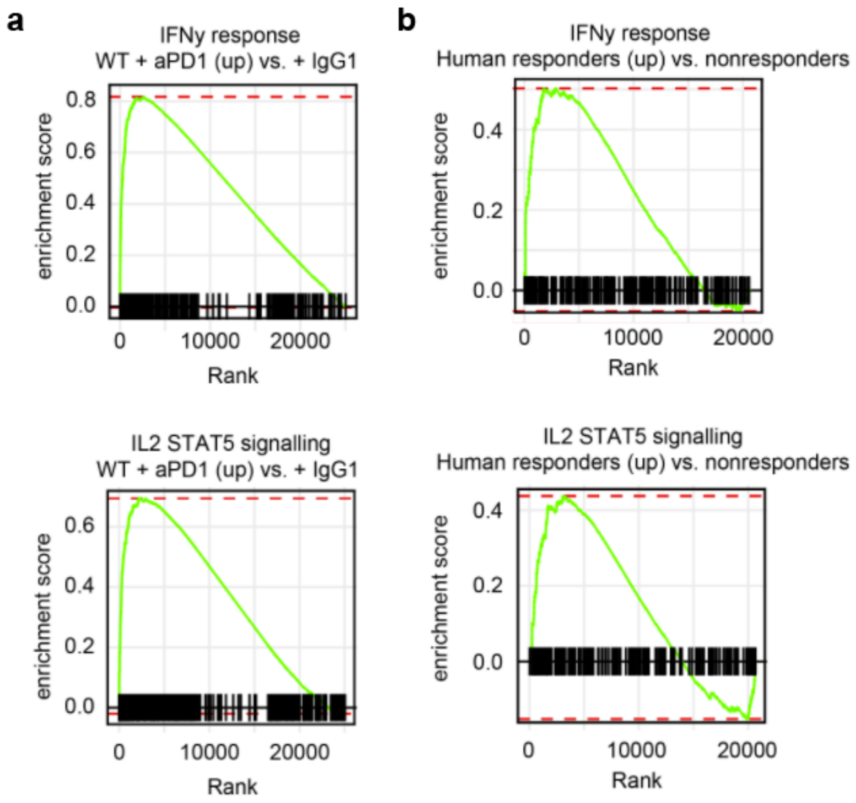


**Supplemental Figure S7 | Gene set enrichment analyses revealing the biological signifcance of ICB response. a**, Enrichment plots from GSEA showing the enrichment in immune pathways (IFNγ response and IL2-STAT5 signaling) of αPD1-treated WT tumors relative to isotype controls (n = 5). **b**, Enrichment plots showing the enrichment in immune pathways of αPD1-treated tumors from responsding (CR + PR) relative to non-responding (PD) patients.

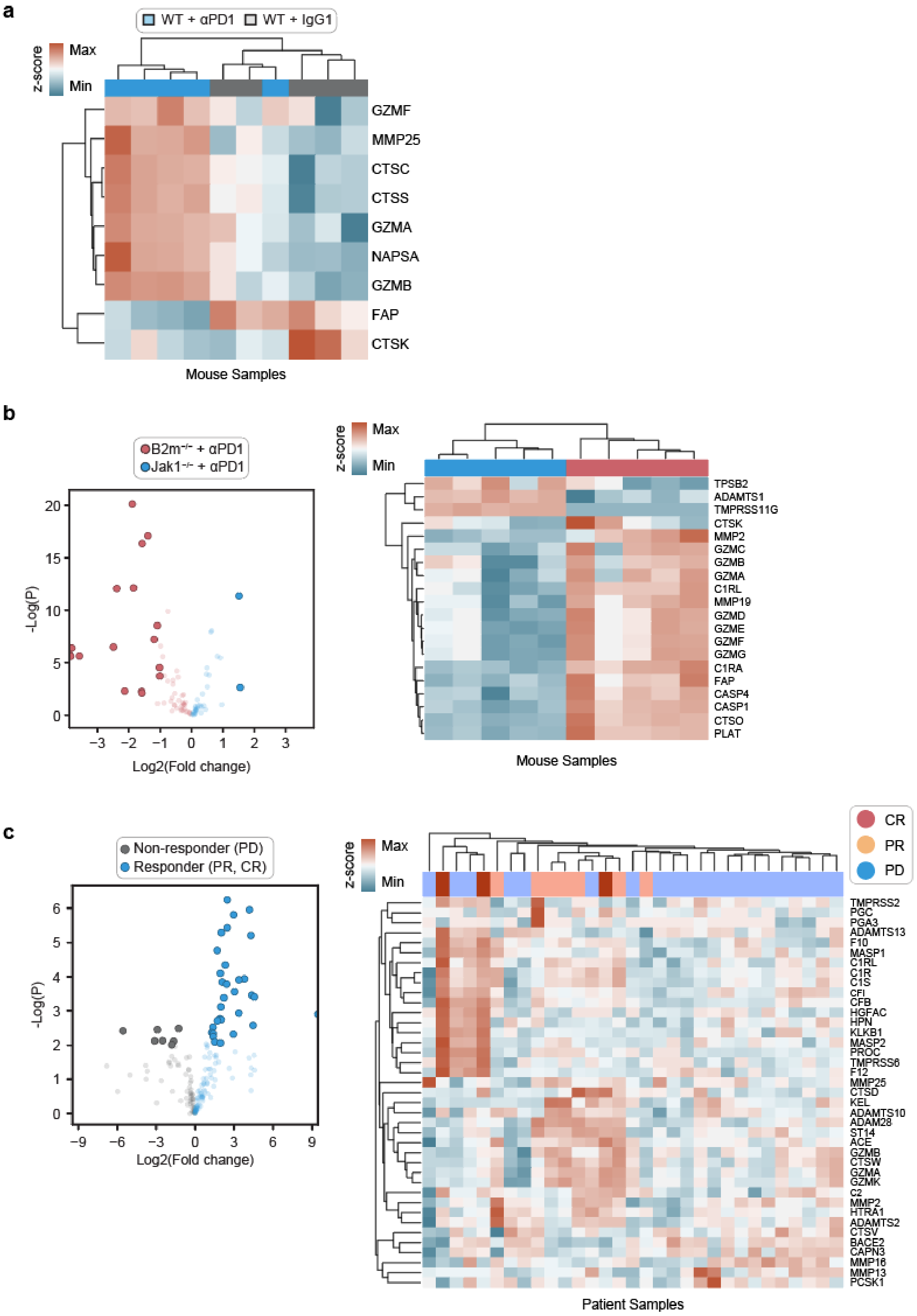

a
WT + αPD1
WT + IgG1
z-score
Max
Min
GZMF
MMP25
CTSC
CTSS
GZMA
NAPSA
GZMB
FAP
CTSK
Mouse Samples
b
B2m−/− + αPD1
Jak1−/− + αPD1
-Log(P)
Log2(Fold change)
z-score
Max
Min
TPSB2
ADAMTS1
TMPRSS11G
CTSK
MMP2
GZMC
GZMB
GZMA
C1RL
MMP19
GZMD
GZME
GZMF
GZMG
C1RA
FAP
CASP4
CASP1
CTSO
PLAT
Mouse Samples
c
Non-responder (PD)
Responder (PR, CR)
-Log(P)
Log2(Fold change)
z-score
Max
Min
CR
PR
PD
TMPRSS2
PGC
PGA3
ADAMTS13
F10
MASP1
C1RL
C1R
C1S
CFI
CFB
HGFAC
HPN
KLKB1
MASP2
PROC
TMPRSS6
F12
MMP25
CTSD
KEL
ADAMTS10
ADAM28
ST14
ACE
GZMB
CTSW
GZMA
GZMK
C2
MMP2
HTRA1
ADAMTS2
CTSV
BACE2
CAPN3
MMP16
MMP13
PCSK1
Patient Samples

**Supplemental Figure S8 | Proteases are differentially expressed in ICB response and resistance. a**, Heatmaps showing row-normalized expression (FPKM) of proteases differentially expressed between αPD1-treated WT tumors and IgG1-treated controls (n = 5). **b**, (Left) Volcano plots summarizing differentially expressed proteases between αPD1-treated B2m$^{-/-}$ and Jak1$^{-/-}$ MC38 tumors (n = 5). The threshold for differentially expressed genes (opaque dots) was defined as P value ≤ 0.05 and |log2(fold change)| ≥ 1. (Right) Heatmaps showing row-normalized expression (FPKM) of proteases differentially expressed between B2m$^{-/-}$ and Jak1$^{-/-}$ MC38 tumors (n = 5). **c**, (Left) Volcano plots summarizing differentially expressed proteases between human tumors from responsders (CR + PR) and non-responders (PD) (n = 5). The threshold for differentially expressed genes was defined as P value ≤ 0.01 and |log2(fold change)| ≥ 1. (Right) Heatmaps showing row-normalized expression (FPKM) of proteases differentially expressed between human tumors from responsders (CR + PR) and non-responders (PD) (n = 5).

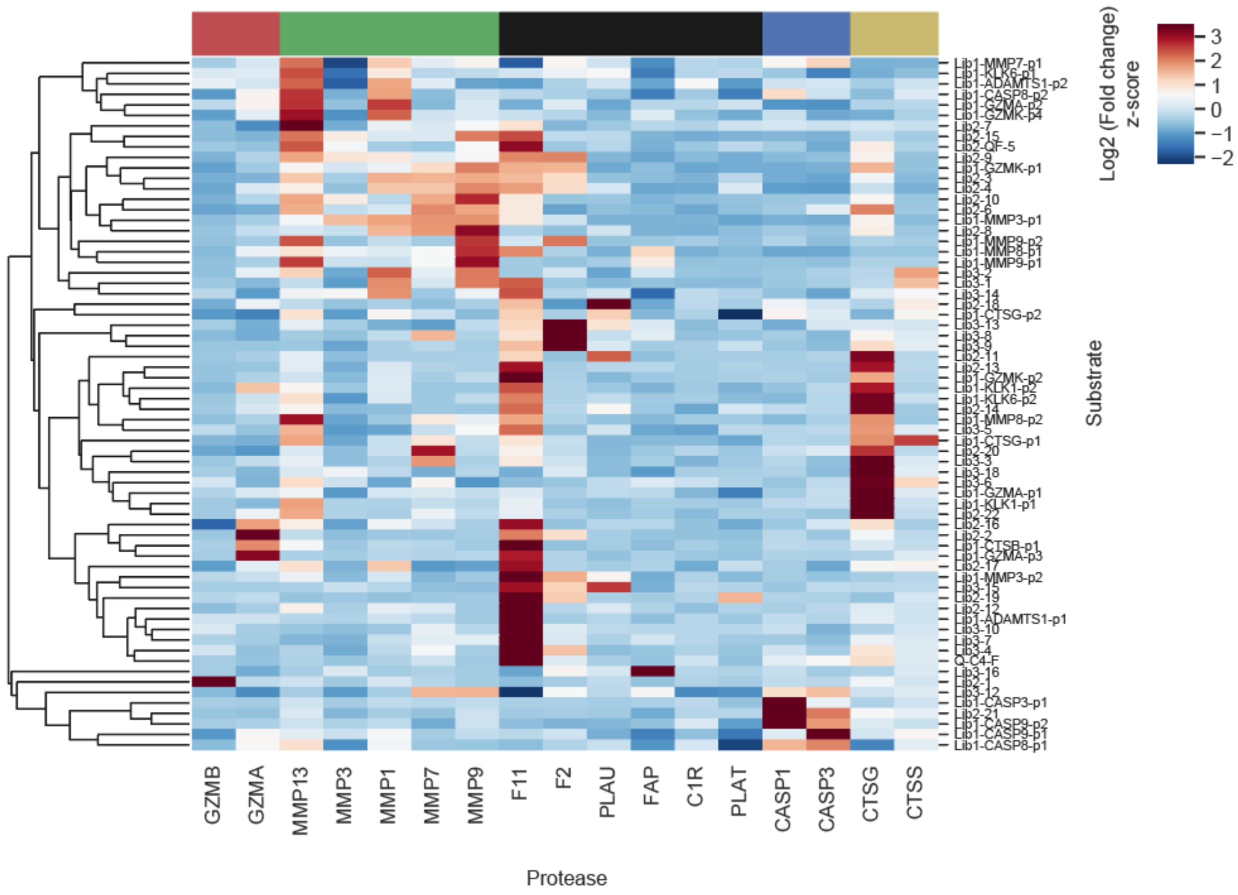


**Supplemental Figure S9 | Optimization of peptide substrates for target proteases.** Heat map summarizing the log2 fold change in fluorescence of 66 quenched substrates at 60 minutes after addition of the respective recombinant protease (n = 3). Signals were row-normalized before plotted.

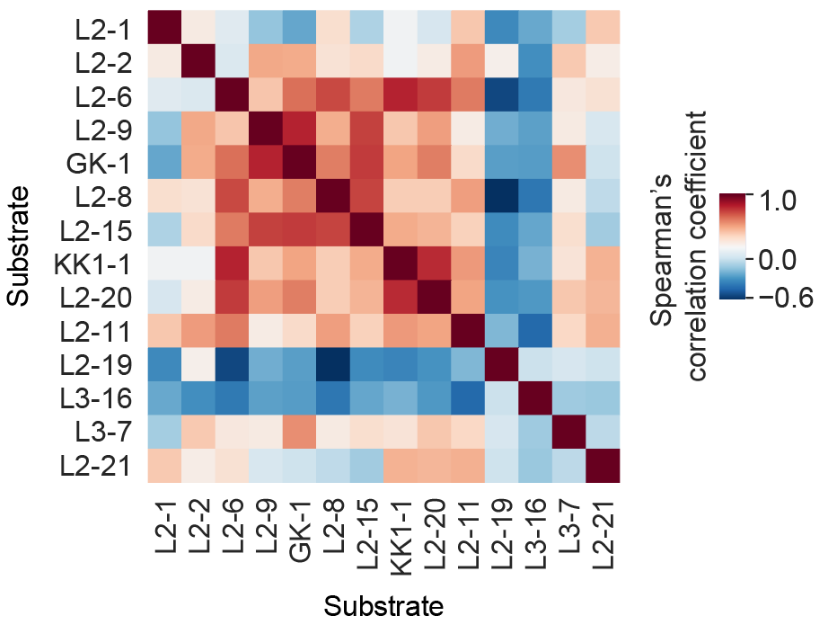


**Supplemental Figure S10 | Correlation analysis of substrate cleavage signatures.** Correlation matrix showing the Spearman's pairwise correlation coefficients between the cleavage signatures of 14 peptide substrates in the INSIGHT panel.

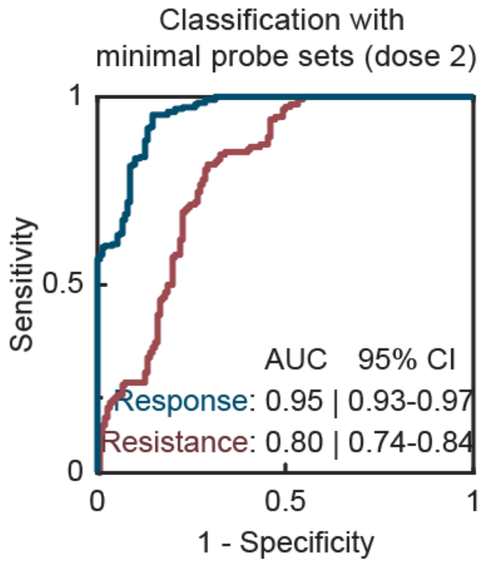


**Supplemental Figure S11 | Classification performance using minimal probe sets based on dose 2 urine signals.** Area under the ROC curve analysis showing the diagnostic specificity and sensitivity of random forest classifiers based on the minimal set of 3 probes (L2-8, L3-7, L2-1) for response monitoring (AUC = 0.95, 95% CI = 0.93-0.97) and on the set of 5 probes (L2-11, L2-20, L2-19, L3-16, and L2-9) for resistance stratification (AUC = 0.80, 95% CI = 0.74-0.84).

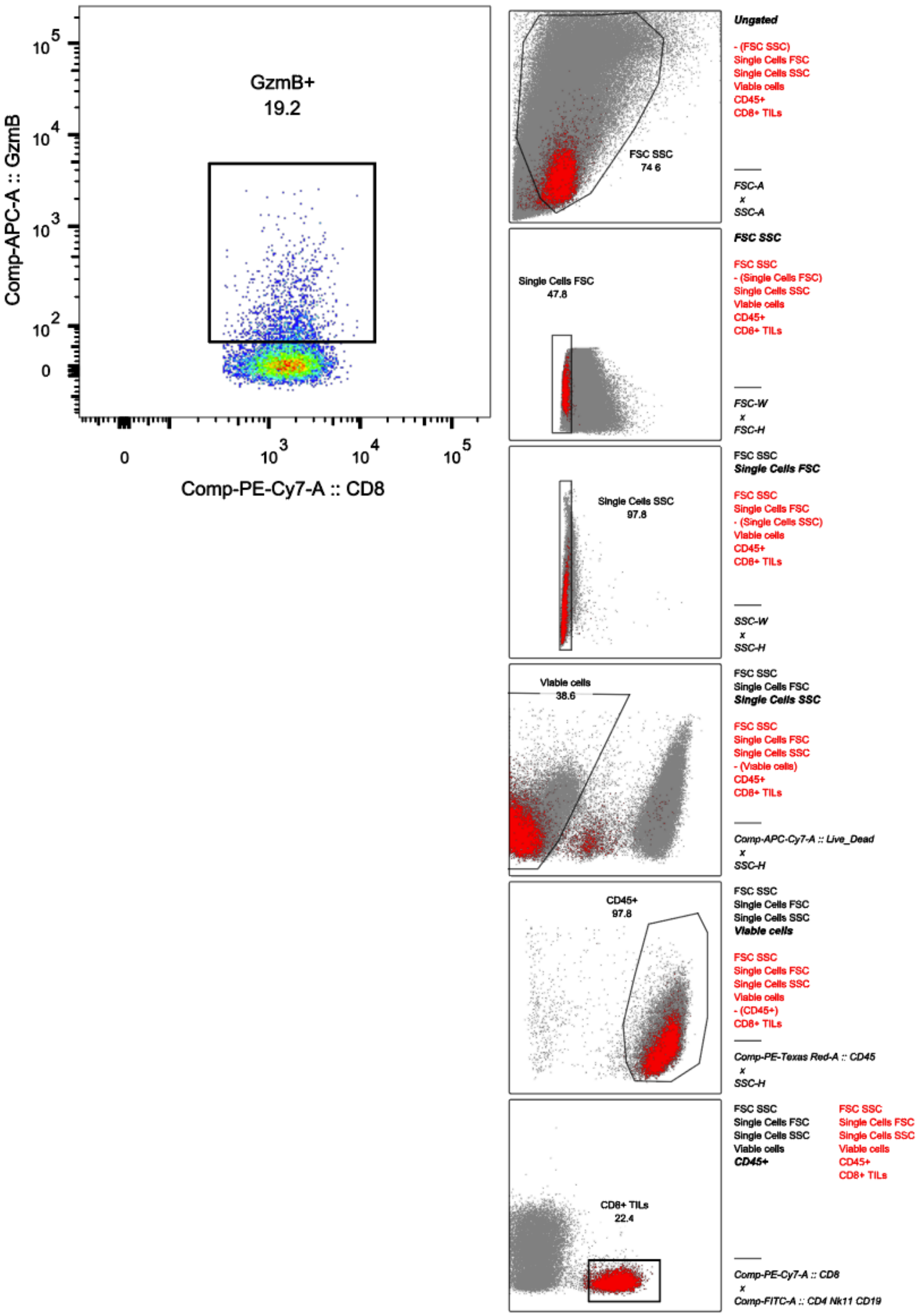


**Supplemental Figure S12 | Gating strategy to identify viable CD8+ TILs expressing GzmB.**
